# Evolution in microbial microcosms is highly parallel regardless of the presence of interacting species

**DOI:** 10.1101/2023.12.14.571477

**Authors:** Nittay Meroz, Tal Livny, Gal Toledano, Yael Sorokin, Nesli Tovi, Jonathan Friedman

**Affiliations:** Institute of Environmental Sciences, Hebrew University, Rehovot, Israel; The Rachel and Selim Benin School of Computer Science and Engineering, Hebrew University, Jerusalem, Israel

## Abstract

During laboratory evolution, replicate bacterial populations often follow similar trajectories, thus their evolution is potentially predictable. However, predicting the evolution of natural populations, which are commonly embedded in multispecies communities, would prove extremely difficult if adaptations are contingent on the identity of the interacting species. The extent to which adaptations typically depend on coevolving partners remains poorly understood, since coevolution is commonly studied using small-scale experiments involving few species, making it challenging to extract general trends. To address this knowledge gap, we study the adaptations that occurred in strains of each of 11 species that were either evolved in monoculture or in multiple pairwise co-cultures. While we detect slight but significant partner-specific effects we find that the majority of evolutionary changes that occur are robust across strains that evolved with different partners; species’ growth abilities increase by a similar factor regardless of partners’ identity, shifts in community compositions and interactions are similar between pairs of coevolved and separately evolved strains, and the majority of parallelly mutated genes were detected in multiple biotic conditions. We hypothesized that these results might arise from the fact that ancestral strains are maladapted to the abiotic environment, thus having a pool of adaptations that are beneficial regardless of the biotic partners. Therefore, we conducted a second experiment with strains that were pre-adapted to the abiotic conditions before being co-cultured. We find that even after ∼400 generations of pre-adaptation, evolution is surprisingly non-partner-specific. Further work is required in order to elucidate the factors that influence partner-specificity of coevolution, but our results suggest that selection imposed by the biotic environment may play a secondary role to that imposed by abiotic conditions, making predictions regarding coevolutionary dynamics less challenging than previously thought.

## Introduction

The degree to which evolution is predictable is a core question in evolutionary biology^1,2^. Evolution experiments involving replicate bacterial populations have demonstrated that evolution can be predictable, as a high degree of parallelism is often observed: mutations in the same genes are repeatedly selected, and similar changes in phenotypes occur across independent populations^3–5^. Naturally, when populations evolve in distinct environments evolutionary outcomes are expected to differ^6^. But it is challenging to anticipate which changes in environmental conditions would cause major shifts in evolution, that would require updating the predictions, and which would lead to only subtle modifications. While it was demonstrated that distinct evolutionary outcomes can emerge even due to subtle changes in the abiotic environment^7–10^, it’s still not clear if differences in the biotic environment would typically cause similar shifts.

Variations in the biotic environment may have evolutionary implications, since they can have pronounced ecological and physiological effects. The presence of an interacting species could lead to the extinction of an otherwise prosperous population^11,12^, to massive shifts in gene expression^13^, and to significant changes in the chemical environment^14,15^. Such effects are expected to change evolutionary outcomes by altering selection pressures^16^ and population sizes^17^, or by creating eco-evolutionary feedback loops^18,19^. Although expectations of this nature are often grounded on strong theoretical foundations^20^, we still lack comprehensive empirical evidence for the typical degree to which evolutionary outcomes depend on the biotic environment.

Coevolution’s capacity to induce notable shifts in evolutionary outcomes was illustrated in a few experimental studies, however, it is still not clear whether these are the rule, or the exception. For example, species lacking the ability to synthesize reciprocal essential metabolites have coevolved to cross-feed^21^; *Bacilus subtilis* that evolved with the black mold fungus *Aspergillus niger* has evolved to better invade the mold’s niche, but did not evolve the same capacity when it evolved without it^22^; and *Escherichia coli* strains that evolved with the predator *Myxococcus xanthus* have acquired a different set of mutations then that of their equivalents that evolved alone^23^. These and other notable examples where evolution was highly partner-specific^24–26^, typically involved a strong and highly defined interaction that is driven mostly by a specific mechanism (with the exception of ref^26^).

However, in experimental systems that studied adaptations in communities of the same guild, biotic-context specific effects are often less pronounced, at least for some of the species involved^27–31^. For example, in a recent study, strains of 5 species that were coevolved did not differ from those that evolved separately in either the growth rates within the community, the assembled community productivity or in invader fitness^29^. Similarly, selection targets of *Stenotrophomonas sp.* strains that were coevolved within a five species polyculture were similar to those that evolved alone, but different from those that were gained when eco-evolutionary feedback was not permitted^30^. Hence, it is still not evident how strongly the presence of an interacting species typically affects evolutionary outcomes.

In order to understand how strongly evolutionary outcomes depend on the presence of specific partners, we study changes that occurred in 11 bacterial species that were either evolved alone (*Monoculture*) or within different co-cultures for ∼400 generations^32^ (Figure 1A). We measure how these strains evolved phenotypically (growth rate, carrying capacity - Figure 1B), and genotypically; and how co-cultures composed of these strains changed in composition and in interspecies interactions (Figure 1B). We then compare the parallelism between strains that evolved in the same evolutionary treatment (same partner or alone, Figure S22B-C), to that of strains that evolved in different evolutionary treatments (different biotic-partners, or alone vs with biotic-partner, Figure S22B-C). We find that parallelism is consistently higher within evolutionary treatments than between treatments, suggesting that the presence of a biotic partner has affected evolutionary outcomes. However, the magnitude of partner-specific effects was generally low - only between 2-5% of the change varied between evolutionary treatments, suggesting that adaptations were only weakly dependent on the biotic context. This was also the case when ecological interactions were strong, and ancestral species were pre-adapted to the abiotic environment. These findings indicate that many predictions based on one set of biotic conditions could remain accurate even when these conditions change.

**Figure 1:**
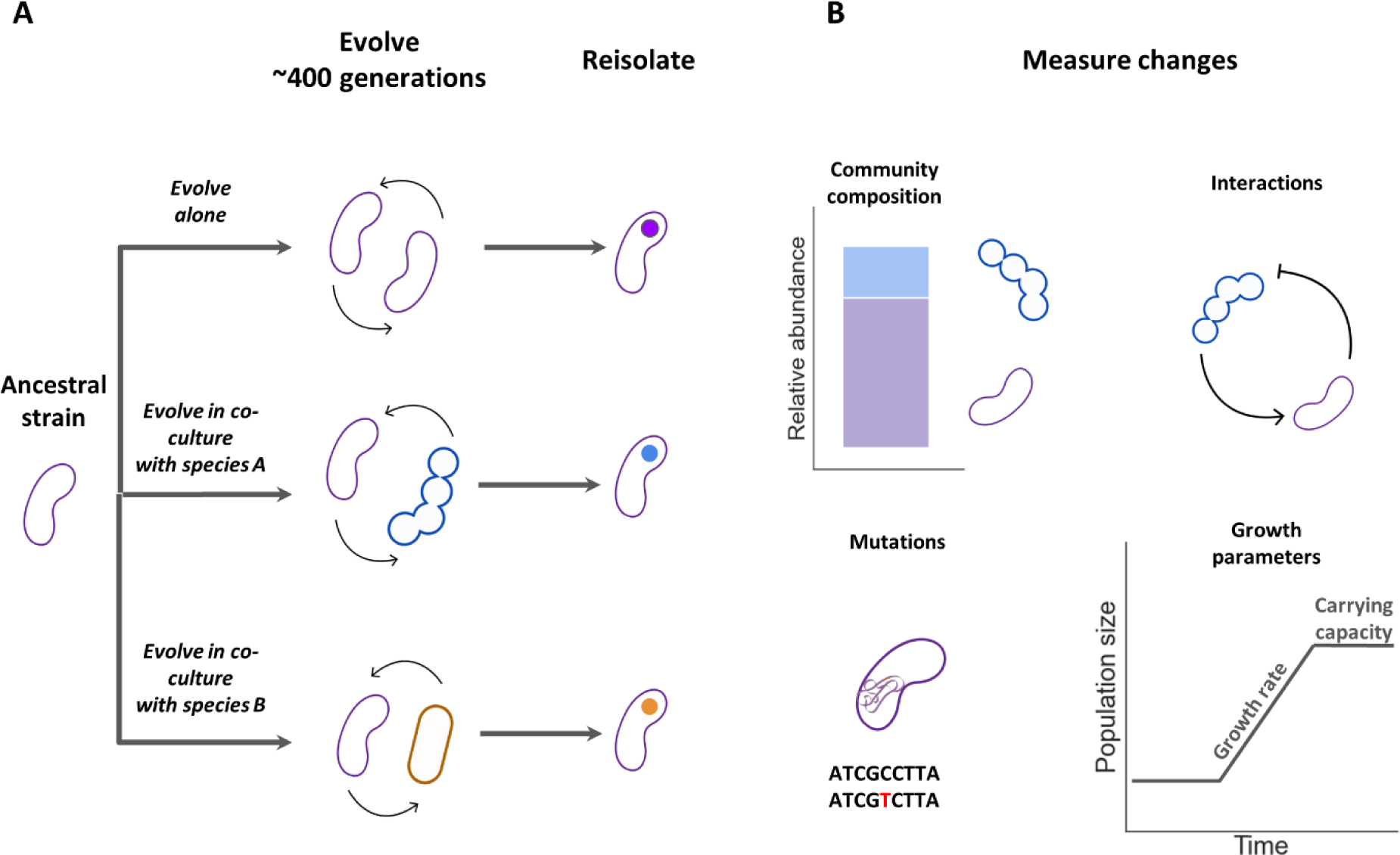
Overview of the experimental design. (A) We evolved ancestral strains of each of 11 species for ∼400 generations either alone (Monoculture), or in different pairwise co-cultures. (B) We measured changes that occurred during the evolution experiment by comparing several parameters of the evolved strains with those of their ancestors: community composition (fractions of the species in co-culture), interactions (growth in co-culture vs growth alone), mutations, growth rate (median per-capita growth rate in exponential phase), and carrying capacity (OD_600_ after 48h growth).

## Results

In order to assess the dependence of evolution on the presence of other species, each of 11 species (Table S1) was evolved alone (monoculture) and in 2-5 different pairwise co-cultures with an average of 4 evolutionary treatments per species (including monoculture and unique pairwise co-cultures, Table S3). Cultures were propagated for ∼400 generations in M9 minimal media supplemented with three carbon sources - acetate, serine, and galacturonic acid^32^ (Methods). After evolution, we reisolated strains and measured the properties of the evolved strains and communities. In most cases (67%) the ancestral species were negatively affected by the presence of their partner, reaching an average 35% 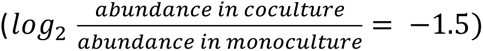 of their monoculture population size in co-culture (Figure S1, S2). In the other 33% of cases, the ancestral species were facilitated by the presence of their partner, reaching on average an abundance of 370%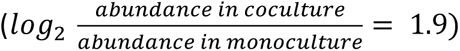 that of their growth alone. Thus, our experimental system includes strains that evolved within a variety of different co-cultures with partners that affected them both positively and negatively.

### Co-culture properties and species traits are mostly shared between evolutionary treatments

The composition of most co-cultures changed significantly during the experiment^32^. Such changes could occur due to species adapting to each other’s presence, or due to each species adapting to the abiotic conditions. In order to distinguish these alternatives, we compared the composition of co-cultures of strains that were coevolved together to the composition of co-cultures of strains that were each evolved separately in monoculture. The changes that occurred in the composition of co-cultures of strains derived from the two evolutionary treatments were strongly correlated (Figure 2A, *Pearson r = 0.8, p = 0.001,* Figure S4), and changes were rarely qualitatively different (Figure 2A : In 12/13 pairs the same species increased in abundance regardless of evolutionary treatment, binomial p-value = 0.003, Figure S4).

**Figure 2:**
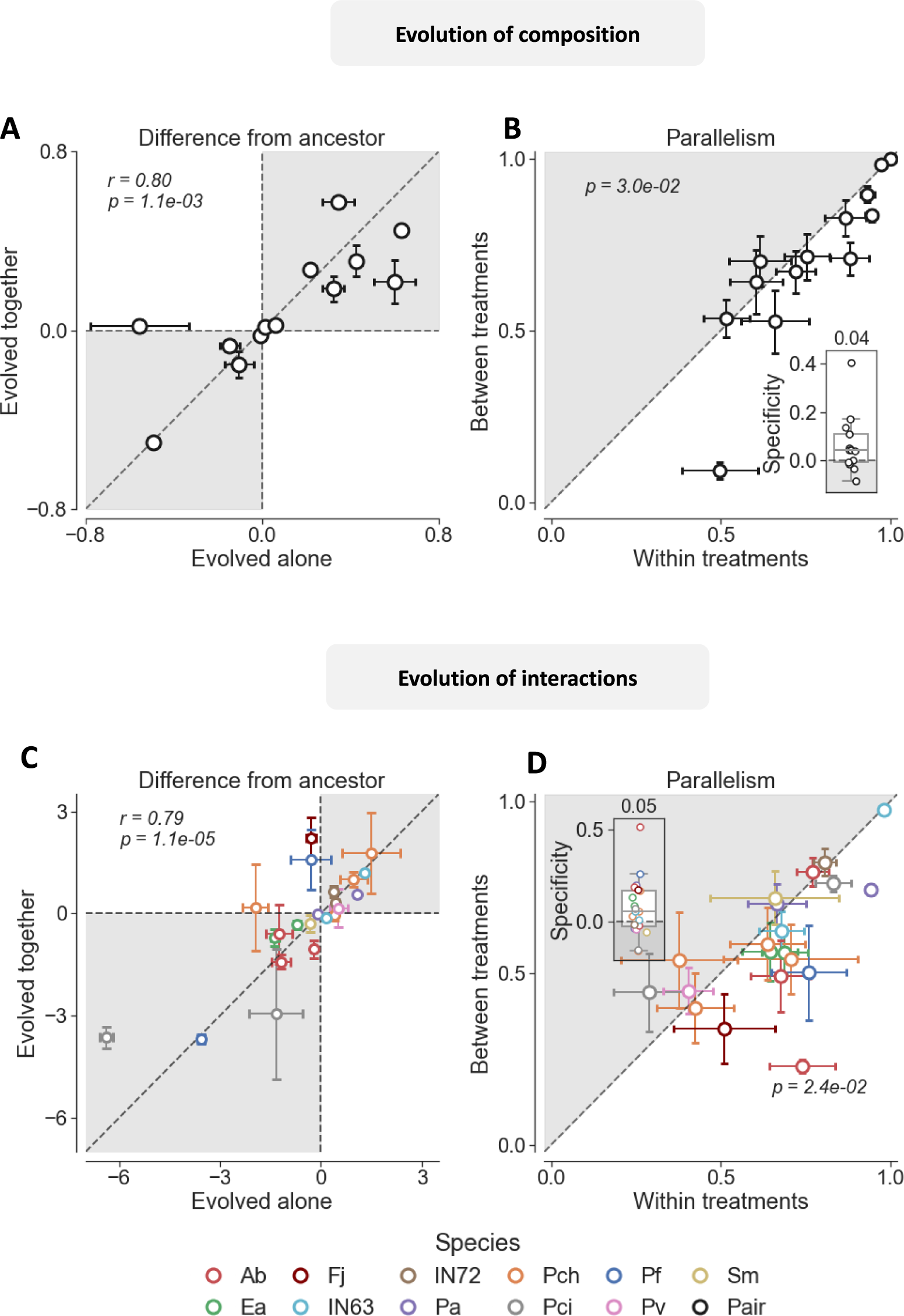
Evolution of co-culture properties is similar regardless of the presence of coevolving species. (A) Change in the composition of pairs of strains that evolved together, against the change in composition after they evolved separately in monocultures. Change in composition is measured as the fraction of a species in the evolved co-culture minus its fraction in the ancestral co-culture. Each circle represents a unique and initially identical pair of species, and the circle centers and error bars represent the mean and the standard error of 2-6 independently evolved co-cultures (Table S4). (B) Parallelism in the evolution of composition within treatment (parallelism between pairs of strains that either evolved alone or coevolved) against the parallelism between treatments (parallelism between pairs strains that evolved alone to strains that coevolved). Circles below the 1:1 line correspond to pairs that did not differ depending on treatment, and circles under the line correspond to ones whose compositional changes were more similar. Circles and error bars indicate the mean and the standard error of each unique pair of species. Insets show the distribution Specificity scores for each parameter, and the number above is the median score across all unique pairs of species. Higher Specificity scores indicate the parallelism within treatments was higher than between treatments. (C) Change in the effect of one species on another’s growth (one-sided interaction) after they evolved together, against the change after evolving separately in monocultures. One-sided interaction is measured as the log_2_ ratio between the abundance of the affected species in co-culture and its abundance when grown alone. Change is quantified as the one-sided interaction in the evolved co-culture minus the one-sided interaction in the ancestral co-culture. Each circle represents the interaction in a unique pair of species, and the circle center and error bars represent the mean and the standard error of 2-5 independently evolved co-cultures (Table S4). Circle color indicates the affected species. (D) Parallelism in the evolution of interactions within treatment (parallelism between pairs of strains that either evolved alone or coevolved) against the parallelism between treatments (parallelism between pairs strains that evolved alone to strains that coevolved). Circles and error bars indicate the mean and the standard error of each unique pair of species. Insets show the distribution specificity scores for each parameter, and the number above is the median score across all unique pairs of species. Text in panels A and B indicates the Pearson r and associated p, text in panels C and D indicate the p-value of a one-sided Wilcoxon test. A, B: Data from experiment Ev1; C, D: data from experiments Ev2-5 (Methods).

In order to quantify the similarity in evolutionary outcomes we devised a measure of parallelism (*Φ*), which corresponds to the fraction of the total amount of evolutionary change in a trait’s value, or a co-culture property, which is shared between independently evolved strains, or pairs of strains (Methods, see Supplementary Information 5 for a detailed information about the calculation). This measure ranges between 0-1, where 0 means that strains evolved in exactly opposing directions and 1 means that trait values are identical between independently evolved strains. Parallelism in the evolution of community composition tended to be higher within treatments than between, suggesting that some shifts in composition were due to species adapting to each other (Figure 2B, one sided paired Wilcoxon test p-value = 0.03). However, parallelism was also high between treatments; the median co-culture shared 0.75 (median *Φ*_*within*_ = 0.75) of the change within treatment and 0.71 of the change between treatments (median *Φ*_*between*_ = 0.71; Figure 2B). We devised a second measure - Specificity score (*Φ*_*within*_ − *Φ*_*between*_), which quantifies the extent to which evolution within treatments is more similar than between treatments (Methods, Supplementary Information 5). Across all pairs, the median Specificity score of the change in community composition was 0.04, indicating that most changes could not be attributed to the effect of species on each other, but was rather due to adaptations to the abiotic environment (Figure 2B inset, combined permutation tests p-value across all pairs for the hypothesis that differences are smaller within groups, p-value = 0.1).

In order to further study how co-culture properties are typically affected by coevolution, we measured changes in species interactions following both coevolution and separate evolution in monoculture. Theoretical studies predict that coevolution can affect species interactions through various mechanisms such as changing the extent of niche overlap, and the intensity of interference competition, or cross-feeding^33–35^, depending on the mechanism of interaction and environmental conditions. However, interactions could evolve also through adaptations to abiotic factors which occur also in the absence of the interaction (e.g. due to better utilization of the supplied nutrients). We find that the strength of interactions (quantified as the log_2_ ratio between a species abundance in a specific co-culture and its abundance when grown alone) evolved significantly in many pairs during the experiment (Fig 2C, Figure S5). However, evolution of interactions seemed mostly non-related to coevolution. Changes in interactions were strongly correlated between pairs of strains that evolved separately and together (Fig 2C, *Pearson r = 0.79, p =* 10^−5^), and the direction of change typically did not differ between them (FIgure 2C, 17/22 of the interactions changed to the same direction regardless of evolutionary treatment, binomial test p-value = 0.01, Figure S5). Furthermore, while parallelism was higher within treatment (one sided paired Wilcoxon test p-value = 0.02, median parallelism *Φ*_*within*_ = 0.67, median *Φ*_*between*_ = 0.58, Figure S5), evolutionary treatment only accounted for 0.05 of the change in interaction (median Specificity score, Figure 2D, combined permutation tests p-value across all pairs for the hypothesis that differences are smaller within groups, p-value = 0.22). Overall, although some of the change in interactions can be linked to species adapting to one another, most of it occurs regardless of whether the interacting strains were coevolved or each evolved separately in monoculture.

Next, we wanted to understand how coevolving with different partners affected the evolution of a species’ ability to grow in monoculture. For this purpose, we measured the growth rate and carrying capacity of multiple strains that evolved in multiple treatments (evolved alone or in co-cultures with different partners). For both parameters, changes that occurred during evolution in monoculture were strongly correlated with those that occurred during coevolution (Figure S6A,B, growth rate: Pearson r = 0.63, p-value = 0.052; carrying capacity: Pearson r = 0.98, p-value = 2 ∗ 10^−7^). Despite this trend, we note a case where evolution of growth abilities was distinct between evolutionary treatments: *Pf*‘s population size increased by a factor of ∼16 when it evolved alone, but rarely showed a similar increase when coevolved (Figure S7, permutation test p-value after Bonferroni correction, p-value = 0.06). This might be related to the fact that the ancestral *Pf* grows poorly alone, and often benefits from the presence of another species (Figure S1, S2), possibly through cross-feeding of essential metabolites; while evolving the ability to produce these nutrients might be favored in monoculture, it might not be as favorable in co-culture, thus maintaining coevolved strains more dependant on their partners. Overall, similar to the trends shown in co-culture properties, parallelism was consistently higher within evolutionary treatment (Figure S6C, one sided paired Wilcoxon test p-value = 0.05), but most of the change in growth parameters is shared between treatments (median Specificity score= 0.02, combined permutation tests p-value across all species for the hypothesis that differences are smaller within groups, p-value = 0.007), suggesting that the evolution of growth abilities was only weakly affected by coevolution.

### Genetic parallelism is high between evolutionary treatments

While the evolution of co-culture properties and growth abilities was mostly shared between strains that underwent different evolutionary treatments, we wanted to understand whether similar trends are seen also at the genomic level. This might hint to whether the similarity in phenotypes arose due to similar selection pressures, or due to other mechanisms. For this purpose, we sequenced the genomes of 143 evolved strains of 6 different species (*Ab, Pa, Pch, Pf, Ea, Sm*) that evolved alone or with different partners and identified mutations (Methods).

Most species had between 1-3 mutations per strain, except *Sm* that had an average of 33.2 (±2.6) mutations per strain, and is likely a hypermutator (Figure 3A). This is supported by the fact that we identified a 6-bp deletion in mutL, a DNA mismatch repair gene, in the ancestral *Sm* strain. In non-hypermutators, strains that evolved in a co-culture tended to accumulate less mutations than strains that evolved alone, supporting the notion that coevolution might constrain evolution^28,36,37^ (Figure 3A, 4/5 species with less mutations on average, one-sided Wilcoxon p-value = 0.06). Non-synonymous SNPs were the most abundant mutation type in 4/6 species (Figure S8), consistent with adaptive evolution. In *Pa* small indels were slightly more abundant than non-synonymous SNPs, and *Ab* accumulated mostly small indels and SNPs in intergenic regions. We also identified large deletions in all species except in *Sm* (Figure S8).

**Figure 3:**
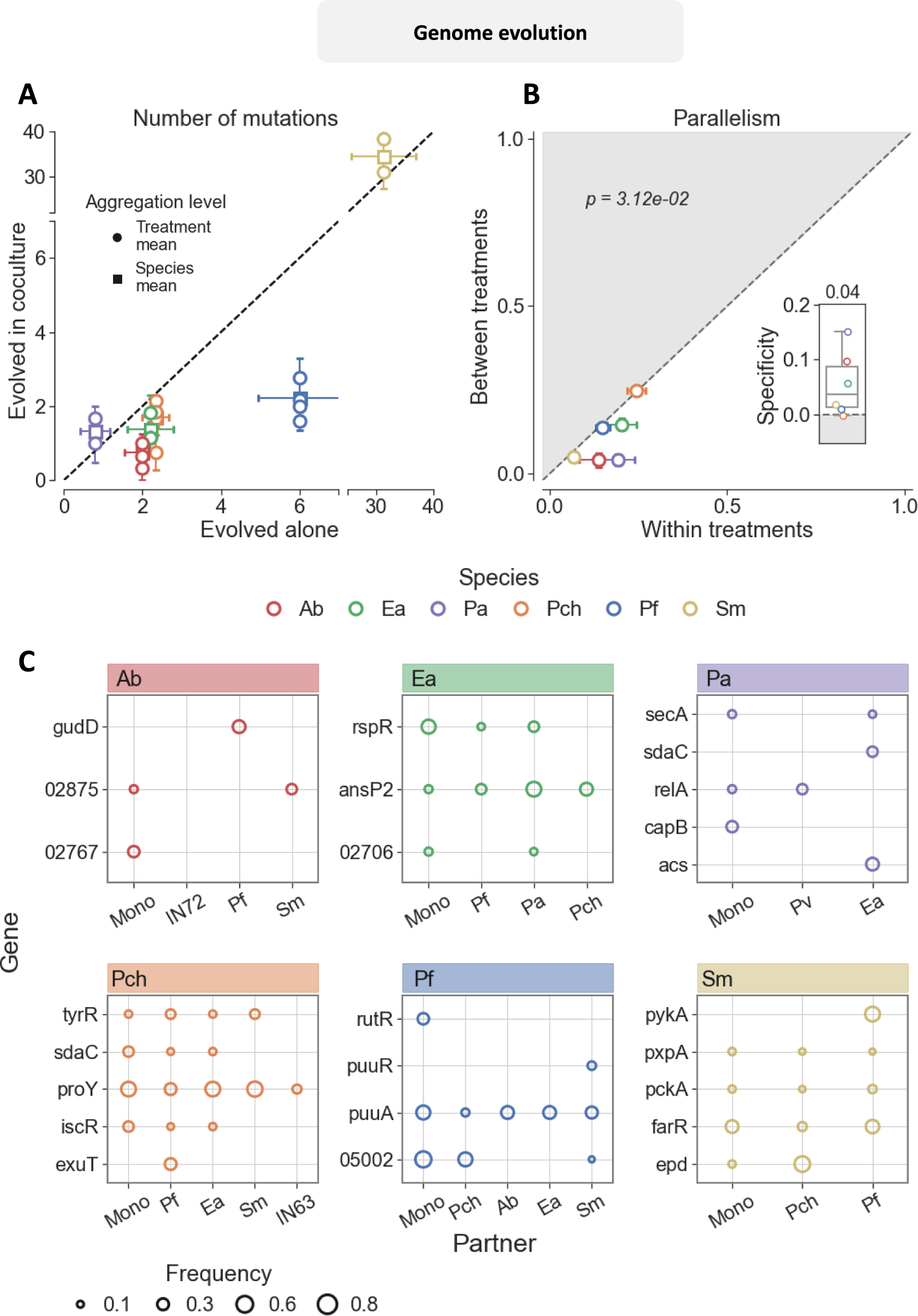
Most parallelly mutated genes are mutated in multiple treatments. (A) Mean number of mutations per strain of species when evolved alone vs when evolved in co-culture. Squares indicate the mean number across all strains of the same species that evolved in co-culture (grand mean), and circles indicate strains that evolved with a specific partner (treatment mean). Error bars denote the standard error of the mean of 3-9 strains (Table S4) (B) Parallelism between mutated genes in strains evolved in the same treatment, against the Parallelism between strains that evolved in different treatments. Parallelism in genomic evolution is measured as the fraction of mutated genes shared between strains (Methods). Error bars indicate the standard error of the mean for each species across 12-31 mutated strains (Table S4). Text indicates the p-value of a one-sided Wilcoxon test. Insets show the distribution Specificity scores and the number above is the median score across all species. (C) Parallelly mutated genes in each species. Genes shown in this plot are the 5% whose observed number of mutations exceeded the expected number of mutations the most (Methods). Colors correspond to species and columns indicate the partner it evolved with (treatment), where ‘Mono’ indicates it evolved alone. Marker size indicates the fraction of strains that evolved in the specific treatment that were mut ated in this gene. Gene names written as a number are hypothetical proteins.

We focus our genomics analysis on gene-level parallelism; For each pair of strains, gene-level parallelism indicates the proportion of genes that were mutated in both strains, relative to the total number of mutations (Equivalent to the commonly used Dice similarity^7^, Methods). Within evolutionary treatment, two strains shared on average between 0.06 (*Sm*, hypermutator) to 0.25 (*Pch*) of the mutated genes (median *Φ*_*within*_ = 0.17, Figure 3B, Figure S9), comparable to other studies (lower than in ref^8^ and similar to ref^7^). Parallelism was consistently lower between treatments (median *Φ*_*between*_ = 0.1, Figure 3B, one sided paired Wilcoxon test p-value = 0.03), suggesting that selection forces varied when species evolved with different partners. However, the median Specificity score across species was only 0.04 (Figure 3B inset, combined permutation tests p-value across all species for the hypothesis that differences are smaller within groups, p-value = 0.003), demonstrating that while partner specific effects were present and detectable most genomic changes were not affected by the presence of another species.

Next, we wanted to identify specific genes that were differently mutated between strains that evolved with a specific partner. To achieve this, we focused solely on genes that were mutated more than expected by chance, which we refer to as parallelly mutated genes (Figure 3C, Methods). 73% of the parallelly mutated genes were mutated in more than one evolutionary treatment (Figure 3C, Figure S10 shows the results applying a different criterion, which accounts for the fact that a bias towards genes that are mutated across treatments could arise), implying that the selection for these mutations was not solely due to the presence or absence of a specific species. However, some genes were mutated predominantly or exclusively in a specific evolutionary treatment (Figure 3C, table S5). For example, *Sm* had non-synonymous mutations in the gene coding for Pyruvate Kinase (pykA) in 6/9 strains that were evolved with *Pf*, but was not mutated in this gene in any of the 14 strains that evolved without *Pf* (Boschloo p-value after Benjamini Hochberg FDR correction = 0.04). Alternatively, mutations in D-erythrose-4-phosphate dehydrogenase (epd), an intermediate in the Calvin cycle and the pentose phosphate pathway, were mutated in 6/8 *Sm* strains that evolved with *Pch* and only in 1/15 in all other treatments (Boschloo p-value after Benjamini Hochberg FDR correction = 0.06). Other candidate partner-specific affected genes are listed in table S4. Overall, while mutations in some genes appear to be contingent on the presence of a specific partner, the majority seem to be linked to adaptations to the abiotic environment

### Pre-adaptation to the abiotic context does not increase partner-specificity

We hypothesized that the strong parallelism observed across evolutionary treatments may arise due to maladaptation of the ancestral strains to the abiotic environment. This could occur as many early adaptations are likely to be less fine-grained and associated with traits unaffected by specific interactions, contributing to high fitness across different treatments^38^. For example, if a high proportion of the available mutations could confer adaptation to the temperature, shaking conditions, or acidity, these would probably be adaptive regardless of the presence of a specific partner. To test this hypothesis we performed a second evolution experiment with strains that were pre-adapted to the experimental conditions (Figure 4A). We re-isolated 11 strains (one from each species) that were evolved alone for ∼400 generations and used these as pre-adapted ancestors in the second experiment (Supplementary Information 1.2). Similar to the first experiment, these pre-adapted ancestors were propagated either as monoculture, or in co-culture for ∼400 generations.

**Figure 4:**
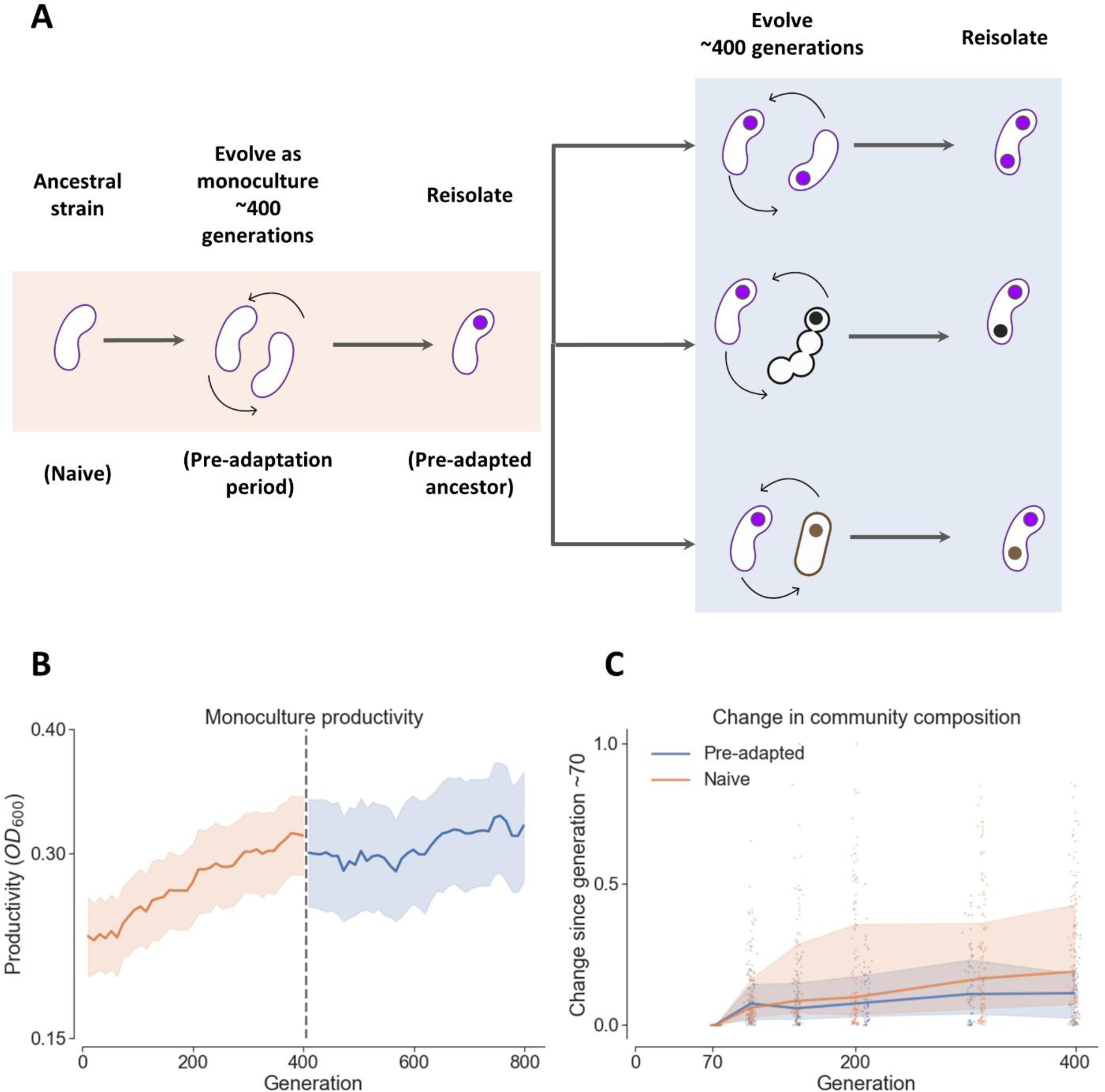
The Amount of change in community composition and growth abilities is reduced significantly in pre-adapted strains and co-cultures. (A) Pre-adapted strains are strains were evolved alone for ∼400 generations and re-isolated before being co-cultured. 17 pairs of pre-adapted strains were co-cultured for an additional ∼400 generations (Methods) (B) Productivity, quantified as the OD_600_ at the end of each growth cycle, of monocultures that evolved in both experiments. The orange area indicates the naive strains, with no prior exposure to the experimental conditions. The blue area indicates pre-adapted strains that were re-isolated at the end of the first experiment. Lines are medians and the shaded area is the interquartile range across all species (C) Change in community composition over evolutionary timescales measured as the Euclidean distance between the composition of each replicate at each time point and its composition at generation ∼70. Generation ∼70 is used here as the proxy for the beginning of evolutionary-, rather than ecological time scales, following previous work^32^. Orange and blue lines and shaded areas denote the median and interquartile range of naive and pre-adapted pairs respectively. B, C: Data from experiments Ev1 (orange), and Ev2 (blue) (Methods).

As expected, the rate of adaptation of pre-adapted strains and co-cultures was lower than that of their naive ancestors. Previous long-term evolution experiments had demonstrated that evolutionary changes in a constant environment could continue for tens of thousands of generations, but that the rate of adaptation decreases with time^39^. Consistent with these findings, in our experiments, growth abilities increased rapidly during the pre-adaptation period, and continued to increase, albeit at a lower pace, during the second evolution experiment (Figure 4B, Figure S18A, B, 7/11 species increased less in growth rate one-sided Wilocoxon p-value = 0.2, 9/11 species increased less in carrying capacity one-sided Wilocoxon p-value = 0.02). Furthermore, pre-adapted strains tended to accumulate less mutations then their naive equivalents (Figure S18E, 4/5 species had less mutations, one-sided Wilocoxon p-value = 0.31). Even though pre-adapted pairs were not previously exposed to their biotic partner, the rate of change in composition was also reduced (Figure 4C, Figure S18C, D, 9/14 pairs changed less in composition, one-sided Wilocoxon p-value = 0.01). This result further supports the notion that most changes that occurred in composition of naive pairs were caused by adaptation to abiotic conditions, and not to the biotic partner. Finally, the decreased evolutionary rates had strengthened our expectations that adaptations towards specific partners would be more pronounced in the pre-adapted pairs.

However, we found that evolution was also highly parallel between treatments after strains were pre-adapted to the abiotic conditions. Evolution after the pre-adaptation period was typically slightly less parallel both within, and between treatments (Figure S19, one sided Wilcoxon test p-value = 0.008, median decrease in parallelism 0.07), suggesting that pre-adapted strains experienced lower selective pressures than their naive counterparts. However, Specificity score did not change significantly in the pre-adapted strains and remained between 0-0.05 (Figure 5, combined Mann-Whitney U-tests within p-value = 0.9, see supplementary materials section 3 for all). These results suggest that the fact that similar evolutionary changes occurred across evolutionary treatment is unlikely to be primarily due to maladaptation of the ancestral strains to the abiotic experimental conditions.

**Figure 5:**
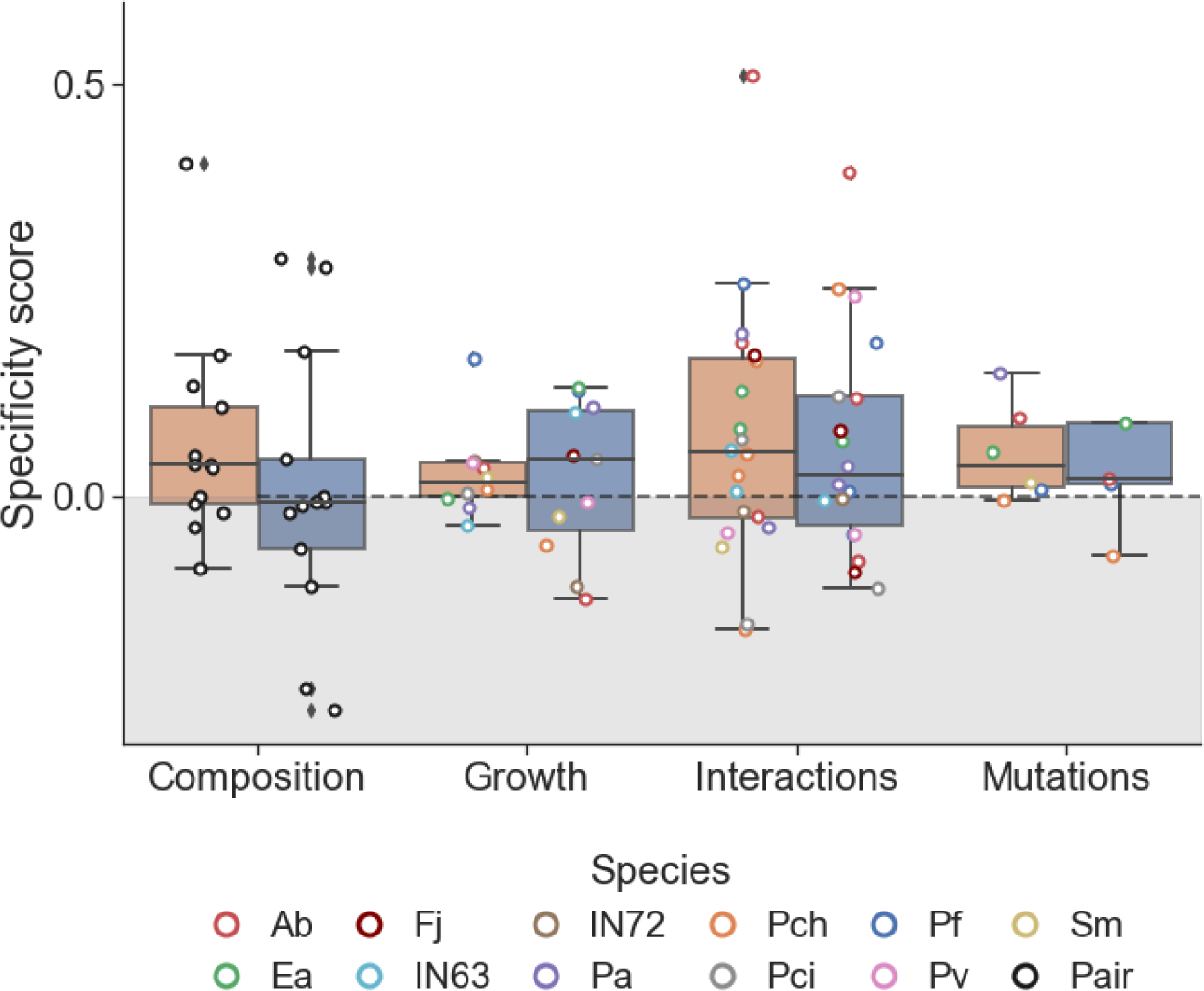
Pre-adaptation does not increase partner specificity. Distribution of Specificity scores for naive (orange boxes) and pre-adapted (blue boxes) strains and co-cultures. Boxes indicate the quartiles and whiskers are expanded to include values no further than 1.5X interquartile range. Circles are specific species (colored) or co-cultures (black). Combined p-value of 4 Mann-Whitney U test for the null hypothesis that naive and pre-adapted strains differ in the specificity score 0.9.

## Discussion

The aim of this study was to empirically test how parallel evolutionary outcomes are when species evolve with or without different biotic partners. We found that while partner-specific effects exist, these typically constitute a relatively small fraction of the overall evolutionary change. These results demonstrate that, at least in some scenarios, evolutionary outcomes could be well predicted without accounting for the effects of specific biotic partners, making predictions less challenging than previously thought. This includes predictions regarding individual species as well as properties of the community. While here we test only pairs of species, we have previously shown that evolutionary changes in composition of trios are typically consistent with those that occur in pairs^32^, suggesting these results may also apply to more diverse communities. However, extending these results to different systems still requires a better understanding of why partner-specific effects were typically weak in our system, and which conditions are expected to promote stronger partner-specific effects.

The degree of similarity in evolutionary outcomes between different conditions likely depends mainly on the similarity of the selective forces imposed by those conditions (Figure 6). In the context of the evolution of communities, we expect the similarity of selective forces to be influenced by the balance between the strength of selection exerted by biotic interactions and by common abiotic factors - the stronger the selection imposed by the biotic interactions, the more evolution will be contingent on the presence of specific species. Indeed, strong evolutionary partner-specific effects have been demonstrated for strongly interacting species, such as predator-prey, mutualistic, or host-pathogen pairs^23,40,41^. Furthermore, the impact of biotic interactions on selection is likely to increase with population density, as higher densities enhance both the frequency of encounters between individuals and their collective influence on the environment. Conversely, we expect evolution to be less partner-specific when biotic interactions are weak, or when species are poorly adapted to their abiotic environment.

**Figure 6:**
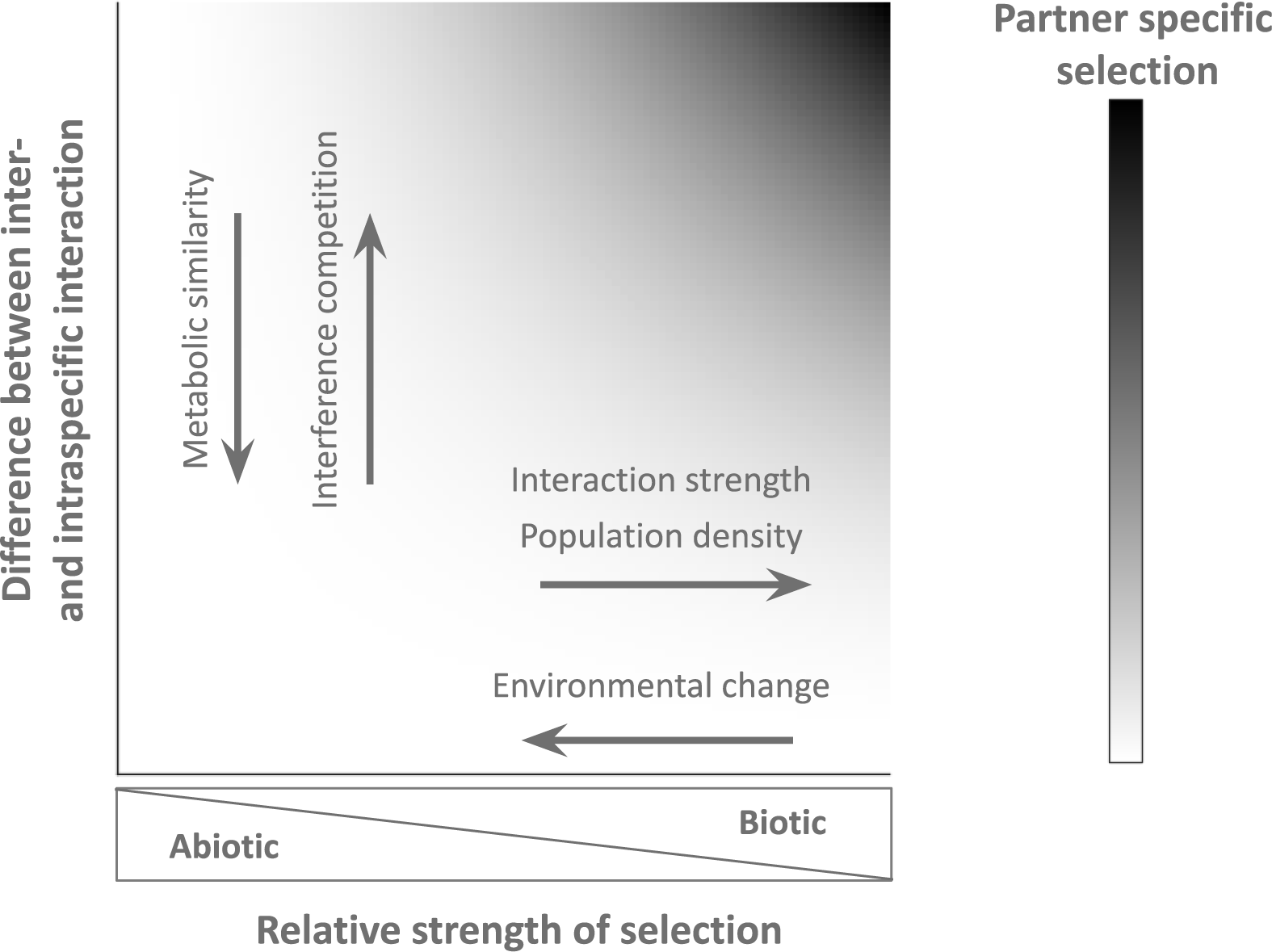
When should species evolution be sensitive to the presence of biotic partners? We propose that the degree to which evolutionary outcomes depend on community composition is contingent on both the strength of biotic selection pressures relative to that of abiotic pressures, and the similarity between inter- and intra-species interactions. Stronger interactions and higher population sizes increase the strength of biotic pressures, while environmental stress increases the strength of abiotic pressures. The similarity between inter- and intra-species interactions is expected to increase with metabolic similarity between the interacting species.

Nevertheless, both adaptation to the abiotic environment and interaction strength are unlikely explanations for the weak partner-specificity of evolution in our system. A ∼400 generations pre-adaptation period did not increase partner-specificity (Figure 5), and some of the strongest interactions resulted in weak or undetectable partner-specific effects. For example, mutated genes of *Pf* strains that evolved with *Ea* were not distinct from those that occurred in *Pf* strains that evolved alone, despite the fact that *Pf* increases its population size by 12-fold when it grows in a co-culture with *Ea (*Figure S2, S9, Figure 3C, Specificity score = 0.0). Furthermore, we find no correlation between interaction-strength to partner-specific effects in any of the measured levels (mutations, evolution of interaction, composition, and growth abilities, Figure S20).

Another factor that can influence the extent to which biotic-partners affect selective forces is the degree of similarity between inter- and intraspecies interactions (Figure 6). If these interactions are similar to each other, individuals within the community may experience selective forces that are largely independent of the presence of specific species even when interspecific interactions are strong. In contrast with predator-prey and host-parasite, where inter- and intraspecific interactions are distinct, species within the same guild can potentially have strong, yet similar strong inter- and intraspecific interactions. For example, if two species utilize resources similarly, and secrete similar metabolites, selection forces experienced by individuals might not differ regardless of their co-occurrence.

However, we also find it unlikely that similar inter- and intraspecific interactions are the main cause of the weak partner-specific effects in our system. Our experiment includes species of different orders (*Pseudomonadales, Enterobacteriales, Flavobacteriales, Mycobacteriales*) which typically have distinct metabolic strategies^42^, and yet, their evolutionary effects on each other were often small. For example, despite the fact that *Enterobacteriales* are commonly acidifiers while *Pseudomonadales* tend to respirate^42,43^, and that *Ea* (*Enterobacteriales*) is strongly inhibited by *Pf* (*Pseudomonadales,* Effect of *Pf on Ea = −1.4*), mutated genes were similar between *Ea* strains that evolved alone and that evolved with *Pf (*Specificity score= −0.0, Figure S2, S9). Under the assumption that phylogenetic distance could serve as a proxy for metabolic dissimilarity, we correlated species phylogenetic distances with the specificity of their evolutionary effects, but found no such correlation (Figure S20). Supporting the notation that strong and distinct interspecies interaction could result in similar selection pressures, a recent study found a strong correlation between the fitness advantage conferred by multiple adaptive mutations in *Saccharomyces cerevisiae* when it grows alone and when the yeast grows with the alga *Chlamydomonas reinhardtii*^44^, despite the fact that these species interacted through obligatory reciprocal nitrogen and carbon exchange^45^.

We speculate that the weak partner-specific effects in our system are mainly due to the growth-dilution dynamics of our experimental system, which could diminish the role of biotic factors in shaping adaptations. Populations in our experiments were propagated using growth-dilution cycles, as commonly done in many evolution experiments^46^. In such a scenario, mutations that arise early in the cycle have a higher fixation probability^47^, and mutations that give benefit at the end of the cycle by increasing yield are not directly under selection ^48^. At the beginning of each cycle, cell-densities are low and selection forces that depend on biotic interaction are likely only significant after cultures reach high densities toward the end of the cycle. However, since most cell divisions were completed by then, beneficial mutations that affect traits related to the interaction could confer a smaller advantage than those that are beneficial earlier in the cycle due to an “early-bird” effect^49,50^. Such dynamics could be further complicated by mechanisms that cause interspecific interactions to change over the course of the growth cycle, such as diauxic growth. Further work is needed in order to understand how growth dynamics affect the partner-specificity of evolution.

It is important to note that distinct evolutionary outcomes could arise even if selection pressures are identical. The presence of another species could impact outcomes by introducing new genetic variation via horizontal gene transfer^51^, or by changing mutation rates^52,53^. The mere change in population size, due to competition or facilitation, might also impact evolutionary outcomes by altering evolutionary rates^17^, or by increasing or decreasing the role of chance in the process^54^.

Finally, several experimental decisions should be taken into account when interpreting our results. First, our experiments were conducted in a well-shaken minimal media provided with three carbon sources (galacturonic acid, acetate, serine). It is possible that a richer growth media, or a spatially structured environment, would produce more eco-evolutionary opportunities that could vary with different partners, thus increasing partner-specificity. Second, our experiments include 11 species of 4 orders, mostly from a single Class (9/11 *Gammaproteobacteria*), and are thus limited in their phylogenetic scope. Additional research is needed in order to understand whether some phylogenetic groups tend to be more evolutionary sensitive to the presence of interacting species, and which mechanisms underlie such bias if it exists. Lastly, our strains evolved for a duration of ∼400 generations per experiment, further work is needed in order to understand whether high parallelism between treatments would be maintained at longer timescales when significant evolutionary changes can accrue. Nevertheless, our results demonstrate that the presence of another species could often have only a marginal effect on evolutionary trajectories, thus suggesting that evolutionary predictions could be less complex than commonly thought.

## Methods

For an easier understanding of the methods used in this work, and considering the data in this study was produced in several separate experiments, we explicitly specify the relevant experiments for each method by noting the relevant experiment numbers at each section, denoted as [’Experiment ID’]. This notation is also used in the figure captions.

### Strains and co-cultures

The set of 11 species used in this study includes environmental isolates and strains from the ATCC collection (Table S1). These species, and all species combinations used in this study are a subset of a larger collection that was used in Meroz at al 2021^32^.

### Growth media

All evolution experiments, co-culturing experiments, and growth assays were conducted in M9 minimal salts media containing 1X M9 salts, 2 mM MgSO4, 0.1 mM CaCl2, 1X trace metal solution (Teknova), supplemented with 3 mM galacturonic acid (Sigma), 6.1 mM Serine (Sigma), and 9.1 mM sodium acetate as carbon sources, which correspond to 16.67 mM carbon atoms for each compound and 50 mM carbon atoms overall.

### Batch culture growth procedure

All evolution experiments and co-culturing experiments were conducted in batch culture with periodic dilutions. In each growth cycle, cultures were grown in 96-well plates (flat bottom) containing 200μl M9 media and were shaken at 900rpm for 48h at 28°C and then diluted by a factor of 1500 into fresh media. OD_600_ was measured at the end of each growth cycle.

### Evolution experiments [Experiments Ev1-2]

Two evolution experiments were conducted following the same protocol. In each experiment cultures were propagated for 38 dilution-growth cycles (see ‘Batch culture growth procedure’), which correspond to ∼400 generations. Community composition was determined in each experiment once every few cycles (see ‘Quantifying community composition’); at transfers 0,2, 5, 7, 10, 14, 19, 30, 38 in the first experiment, and at transfers 0, 4, 7, 10, 14, 20, 29, 39 at the second experiment. At the end of the experiments cultures were mixed with 50% glycerol and frozen at −80°C in 96-deep well plates.

The first experiment (Experiment Ev1) was conducted with strains that were not previously exposed to the experimental conditions, and are thus regarded as ‘naive’ strains. Data from this experiment was previously published^32^, and included 5 species and 21 unique co-cultures were not analyzed in this study due to technical considerations. The second experiment (Experiment Ev2) was initiated with 11 strains (one of each species) that evolved alone in the first experiment. To initiate both experiments, frozen strains were streaked on NB agar plates, and single colonies were inoculated into falcon tubes containing 3ml nutrient broth. After 24h growth, cultures were diluted to OD_600_=10^−2^, and co-cultures were mixed in equal volumes.

We added two naive pairs (Ea-Pa and Pci-IN63), identical to those used in the first experiment to the second experiment in order to verify that the two experiments are comparable. These produced very similar trajectories across the two experiments, therefore verifying that differences between the experiments are due to strains evolutionary history, and not due to technical issues (Figure S25).

### Re-isolations

In order to study how strains changed in growth abilities and in species interactions during the evolution experiments, we re-isolated strains that evolved in co-culture, and strains that evolved alone. While re-isolating coevolved strains was necessary in order to study the species separately, re-isolating strains that evolved alone was done so these would be comparable to the coevolved strains. For re-isolations, frozen stocks of the ∼400 generations-evolved cultures were inoculated into 96-deepwell plates (1ml, Thermo-scientific #260251) containing 500μl M9 using a sterilized 96-pin replicator and incubated at 28°C, shaken at 900rpm. After 48h cultures were diluted and plated on Nutrient Agar plates (5 g/L peptone BD difco, BD Bioscience; 3 g/L yeast extract BD difco, BD Bioscience, 15 g/L agar Bacto, BD Bioscience). Plates were left at room temperature until colonies were detectable and distinguishable (2-4 days). For each strain, 4-8 colonies were picked using a sterile loop, and streaked separately on agar plates to confirm isolation. After colonies were visible, a single colony of each of the 4-8 re-streaked colonies were pooled together in 500μl Nutrient Broth in 96 deep-well plates and grew for 24h at 28°C shaken at 900 rpm. Re-isolated strains were then mixed with 50% glycerol and kept at - 80°C.

### Short ecological experiment for determining the composition of evolved co-cultures [Experiment Ec1]

Community composition was defined as the fraction of species in a co-culture after ∼50 generations of co-culturing. This experiment included two evolutionary treatments for each co-culture: i. co-culture composed of strains evolved separately, ii. co-culture composed of strains evolved together. We used 2-5 independently evolved co-cultures for each evolutionary treatment (evolutionary replicates, Table S3). That is, co-cultures that were identical in the beginning of the evolution experiment, but evolved in different wells for ∼400 generations. Each evolutionary replicate was grown in 3 technical replicates for the duration of this ∼50 generation experiment.

In order to initiate this experiment we inoculated coevolved co-cultures and separately evolved monocultures into 96-deepwell plates containing 500μl M9 using a sterilized 96-pin replicator. Note that these were innoculated from frozen stocks prepared directly from the evolution experiment (Experiment Ev1), rather than from reisolated strains. Plates were incubated at 28°C and were shaken at 900 rpm. After 24h, separately evolved monocultures were co-inoculated by taking 100μl of each species and mixing. Both coevolved and separately evolved co-cultures were diluted by a factor of 1000, and were subjected to 5 dilution-growth cycles. In addition to determining the composition at the end of the experiment (see ‘Quantifying community composition’), it was also determined right after the co-inoculation in order to confirm the presence of both species. If one of the species did not appear in both measurements, the culture was removed from further analysis. The compositions of coevolved pairs in this experiment (Experiment Ec1) correlated well with the compositions in the evolutionary experiment (Experiment Ev1, Ev2) at generation ∼400, suggesting composition is heritable and reproducible (Pearson r = 0.9, p-value = 10^−11^, Figure S26).

### Short ecological experiments for determining the interactions of evolved co-cultures [Experiments Ec2-5]

We quantify the effect one species has on another species growth measured as the log_2_ ratio of the abundance of a species in co-culture and its abundance when grown alone 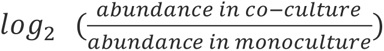. While interactions are composed of the reciprocal effect species have on each other, we analyze the effects separately and refere to these effects as interactions for simplicity. To measure interactions we grew co-cultures and monocultures for 5 dilution-growth cycles, and measured the composition (see ‘Quantifying community composition’) of the co-cultures and OD_600_ of monocultures. The abundance of a species was then quantified as its fraction multiplied by its OD600. For each pair, 1-5 evolutionary replicates were tested (Table S3), i.e., different replicates of the same ancestral strains that underwent the same evolutionary treatment. For each evolutionary replicate, three technical replicates were used.

Isolated strains were transferred using a 96-pin replicator directly from their frozen glycerol stocks into a 96-well plate containing 200 μL nutrient broth and were propagated for 24 hours at 28 °C shaken at 900 rpm. The pairs were then mixed at equal volumes and all cultures were diluted by 10^−3^ into 200 μL of the M9 media, in 96-well plates.

Four separate experiments were conducted in order to measure interactions in this study, since measuring such a large number of interactions in a single experiment is technically challenging. Some interactions were assayed in multiple experiments to check for consistency across experiments, but we only use data from one experiment for each unique pair of species. This was done in order to avoid a bias that can emerge due to differences between experiments. We chose which experiment to use for each pair based on two criteria: (i) The experiment includes all treatments for the pair (ancestor, coevolved, evolved separately); (ii) If more than one experiment includes all treatments, we chose the experiment with the higher total number of evolutionary replicates. The interactions of ancestral strains that were measured in these Experiments (Ec2-5) were correlated with their interactions extracted from the evolution experiments (Ev1-2), demonstrating that interactions are reproducible (Pearson r = 0.86, p = 7*10^−13^, Figure S28).

### Quantifying community composition

Community composition was measured during the evolution experiments and in co-culturing experiments (Experiments Ec1-5) to determine composition and interactions of evolved pairs. Composition was determined by plating and counting colonies, which were distinct in morphology for each species^32^. For that, cultures were diluted in 0.86g/L NaCl solution, by a factor of between 10^7^-2.5*10^8^ and 100μl were plated on Nutrient Agar plates and spread using glass beads. The exact dilution factor varied between experiments in order to reach a large but countable number of colonies of between 20-200. Plates were incubated at room temperature for two-three days and at least 20 colonies were counted manually.

### Quantifying growth parameters [Experiment Gr1]

Isolated strains were transferred using a 96-pin replicator directly from their frozen glycerol stocks into a 96-well plate containing 200μL M9 media and were incubated at 28°C shaken at 900 rpm. After 48h, cultures were diluted by a factor of 1000 to fresh media, and split into 2 technical replicates each, such that technical replicates grow in separate plates. The optical density was then measured using 6 automated plate readers (3 Epoch2 microplate reader - BioTek and 3 Synergy microplate reader - BioTek) simultaneously. Plates were incubated at 28 °C with a 1 °C gradient to avoid condensation on the lid, and were shaken at 250 cpm. OD_600_ was measured every 5 min.

Growth curves were smoothed by a moving average with a window of 50 minutes. Exponential growth phases were determined manually by inspecting each growth curve separately. Growth rates were quantified by calculating the median log_2_ difference between sequential measurements within the exponential growth phase. Carrying capacities were defined as the optical density after 48h of growth. Carrying capacities of re-isolated strains that evolved in monocultures (Experiment Gr1) are correlated with these strains’ optical densities during the evolutionary experiments (Figure S27; Experiments Ev1-2, Pearson r = 0.9, p-value = 5*10^−22^), demonstrating growth abilities are reproducible.

### DNA extraction and genome sequencing

We sequenced genomes of strains of 6 species in order to identify mutations that arose during the experiments. We chose these specific species since they had the best annotated genomes. The ancestors (both naive and pre-adapted) were sequenced using a combination of long-reads (Nanopore) and short reads (Ilumina), and reads were assembled and used as reference genomes. Evolved strains were sequenced using short-reads only.

A single colony of each strain was picked from NB agar plates and incubated overnight at 28°C in 3ml NB. Genomic DNA was extracted from each sample with the Bacterial Genomic DNA Kit (NORGEN Biotek, #17900) according to the manufacturer’s instructions. The libraries for Illumina sequencing were constructed according to standard protocols using the Illumina DNA prep kit and IDT 10 bp UDI indices (Illumina, San Diego, CA, USA) by the sequencing facility at SEQCENTER (PA, USA, https://www.seqcenter.com/). The samples were sequenced on an Illumina NextSeq 2000. Demultiplexing, quality control, and adapter trimming were performed by the sequencing center with bcl-convert. Average coverage for short reads is 121 with a standard deviation of 40. Nanopore samples (long reads) were prepared for sequencing using Oxford Nanopore’s “Genomic DNA by Ligation” kit (SQK-LSK109) and protocol. All samples were run on Nanopore R9 flow cells (R9.4.1) on a MinION. Hybrid assembly of the Illumina and Nanopore reads was performed with Unicycler and assembly annotation was performed with Prokka. Both assembly and annotation of the genome were performed by the sequencing center.

### Mutations Calling

Sequences were trimmed using *Trimmomatic*^55^ (version 0.39), using a sliding-window approach. Reads were clipped when the average quality score was <20 in a 5-20-bp window and to a minimum length of 25 bp. Mutations were identified by comparing evolved strains to their ancestors using *breseq* v. 0.36.1^56^ with default parameters.

Mutations that appeared in more than one independent evolving populations in exactly the same position and had the exact same nucleotide change (SNPs or indels) were inspected manually to avoid a bias that could occur due to limitations of the sequencing procedure and variant calling algorithm. If we found a reason to suspect that this mutation was incorrectly assigned it was filtered out. In Table S7, we list each exact sequence mutation, describing our decision on whether to filter it out and providing the rationale behind our choice. Importantly, we demonstrate that our key results remain qualitatively consistent with different mutation-filtering choices (Figure S30)

### Quantifying parallelism

For each strain trait (e.g. growth rate or fraction in the community), the degree of parallelism (*Φ*_*ij*_) between two strains (i, j) that share a common ancestor is calculated as:

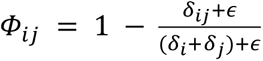

where *δ*_*i*_, *δ*_*j*_ are the Euclidean distances of the trait values of strains i and j from their common ancestor, *δ*_*ij*_ is the Euclidean distance between the two strains, and *ε* is the estimated measurement error of this trait. For each trait, *ε* is calculated as the standard error of the mean of the measured trait values across technical replicates. The Specificity score is defined as the mean difference between parallelism within groups, to the parallelism between groups <u>*Φ*</u>_<u>*within*</u>_ − <u>*Φ*</u>_<u>*between*</u>_. Additional information regarding the measures of parallelism and specificity could be found in Supplementary Information 5.

### Gene-level parallelism

We quantify the gene-level parallelism between independently evolved strains as a measure for their similarity in molecular evolution. Parallelism is quantified in the same manner as parallelism in traits or co-culture properties 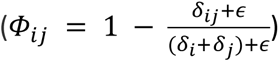, albeit distances (*δ*_*i*_, *δ*_*j*_, *δ*_*ij*_) are measured as Hamming distances rather then Euclidean, and *∈* is not estimated and set to 0. Therefore, *δ*_*i*_ and *δ*_*j*_ are the number of genes mutated in strains *i* and *j*, and *δ*_*ij*_ is the number of genes that were mutated in one of the strains but not in the other. This measure of parallelism is equivalent to the Dice similarity coefficient which is commonly used as a measure of gene-level parallelism^7^ (Supplementary Information 5). In this analysis we exclude synonyms SNP’s and mutations in intergenic regions further than 150 bp upstream to any gene, following other studies^7,57^.

Many of the strains in our dataset include large deletions that affect multiple genes (Figure S7, S15), and might hold adaptive information. To avoid losing this information, we include large deletions in gene-level parallelism calculations. However, these are used conservatively such that each mutational event is included only once even if it affected multiple genes. For example, if two strains include a large deletion where the same two genes were deleted, these would be counted once and would receive the same score as two strains that had one mutated gene in common. Similarly, if one strain had a point mutation in two genes, and another a deletion that included both genes, these would be regarded as one shared mutation. Importantly, the qualitative results remain similar even when multiple-gene deletions are excluded from the analysis (Figure S29).

### Parallelly mutated genes

We identify parallelly-mutated genes by calculating the G-score for goodness of fit between the observed and expected number of mutations for each gene, following ref^58^. Mutated genes include genes that had a non-synonymous SNP, an indel, were included in a large deletion, or were 150 downstream to a intergenic mutation. The expected number of mutations in a gene was calculated as 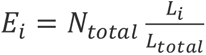, were *L_i_* is length of the gene, *L*_*total*_ is the length of the genome, and *N*_*total*_ is the number of mutations in all strains of a species. The G-score is then calculated as 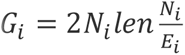. Parallelly mutated genes were then defined as the genes with the 5% largest G-scores (exceeding 20.7).

### Handling technical replicates

Three technical replicates were used for measurements of interactions (Experiments Ec2-5) and community composition (Experiment Ec1), and two technical replicates were used for growth measurements (Experiments Gr1). In each of these cases, technical replicates correspond to strains or co-cultures that evolved in a specific well in the evolutionary experiments (Experiments Ev1-2), and were replicated and split into separate wells in subsequent experiments. Technical replicates were averaged before subsequent analysis, and estimates of the mean and standard errors were calculated using these averaged values, such that they were based on independently evolved strains or co-cultures.

### Permutation tests

We conducted permutation tests for the hypothesis that parallelism is higher within treatments than between treatments for each of the measured features (composition, interactions, growth, mutations). For that, we used the Specificity score <u>*Φ*</u>_<u>*within*</u>_ − <u>*Φ*</u>_<u>*between*</u>_ as the test statistic for each species, interaction, or co-culture. Distinct permutations were generated by sampling the treatment labels without replacement, while ensuring that each label permutation is only sampled once. We included all distinct permutations if there were less than 2,000 such permutations, or 2,000 randomly selected permutations if there were more. For each permutation, we calculated the Specificity score, and p-values are calculated as 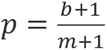, where *b* is the number of permutations that yield a Specificity score greater or equal to the original score, and m is the number of sampled permutations^59^. We subsequently applied a Bonferroni correction to account for multiple tests when needed.

