## Supplementary Information for "Evolution in microbial microcosms is highly parallel regardless of the presence of interacting species"

##### Table of content:

1. Experimental setup and dataset description
    1.1 Experimental setup and dataset description - naive strains
    1.2 Experimental setup and dataset description - pre-adapted strains
2. Supplementary data for naive strains and pairs
3. Supplementary data for pre-adapted strains and pairs
4. Comparison between naive and pre-adapted strains
5. Parallelism quantification
6. Reproducibility
7. Mutations filtering

### 1. Experimental setup and dataset description

| Short Name | Species | source |
|---|---|---|
| Ea | *Enterobacter aerogenes ATCC 13048* | ATCC |
| Pa | *Pseudomonas aurantiaca ATCC 33663* | ATCC |
| Pci | *Pseudomonas citronellolis ATCC 13674* | ATCC |
| Pv | *Pseudomonas veronii ATCC 700474* | ATCC |
| Pch | *Pseudomonas chlororaphis ATCC 9446* | ATCC |
| Pf | *Pseudomonas Fluroescens ATCC 506* | ATCC |
| Sm | *Serratia marcescens ATCC 13880* | ATCC |
| Ab | *Acinetobacter baylyi ATCC 3330* | ATCC |
| Fj | *Flavobacterium johnsonia strain UW101* | ATCC variant |
| IN63 | *Rhodococcus soli* | Wheat plot |
| IN72 | *Pseudomonas sp. BSP5* | Wheat plot |

***Table S1. Species used in this study.*** *Species in this list are used both as the focal species (the species being studied), and as the biotic partners of other species. All species were included in our previous study*[1].

| Short Name | Species | source |
|---|---|---|
| H77 | *Arthrobacter phenanthrenivorans* | Tomato pot |
| H79 | *Delftia lacustris* | Tomato pot |
| H82 | *Pseudomonas putida* | Tomato pot |
| H97 | *Pseudomonas pseudoalcaligenes* | Tomato pot |
| IN65 | *Pseudomonas alcaligenes* | Wheat plot |

***Table S2. Species used in this study as biotic partners.*** *Species in this list are not studied directly, but are used as the biotic partners of other species. All species were included in our previous study*[1].

#### 1.1 Experimental setup and dataset description - naive strains

| Species | Growth<br>Number of independent populations evolved with each species | Mutations<br>Number of independent populations evolved with each species | Interactions<br>Number of independent evolved cocultures evolved in each condition | Composition<br>Number of independent evolved cocultures evolved in each condition |
|---|---|---|---|---|
| Pf | **With reps**<br>Ea 5<br>Pch 3<br>Pf 3<br>Ab 8<br>IN72 4<br>Sm 5 | **With reps**<br>Ea 6<br>Pch 5<br>Pf 5<br>Ab 8<br>Sm 9 | **seprate together**<br>Pch: 3 2<br>Ab: 3 3<br>IN72: 2 2 | **seprate together**<br>Ea_Pf: 3 3<br>Pf_Ab: 2 5<br>Pf_Fj: 2 2<br>Pv_Pf: 2 3 |
| Pch | **With reps**<br>Ea 5<br>Pch 3<br>IN63 4<br>Pci 2<br>Pf 4<br>Sm 5 | **With reps**<br>Ea 6<br>Pch 6<br>IN63 4<br>Pf 7<br>Sm 7 | **seprate together**<br>Ea: 3 4<br>IN63: 3 4<br>Pci: 3 3<br>Pf: 3 2<br>Sm: 2 2 | **seprate together**<br>Ea_Pch: 3 3<br>Pch_IN63: 3 3<br>Pch_Pci: 2 2<br>Pch_Sm: 3 6 |
| Ea | **With reps**<br>Ea 2<br>Pa 5<br>Pch 5<br>Pf 4<br>Pv 3 | **With reps**<br>Ea 5<br>Pa 6<br>Pch 6<br>Pf 6 | **seprate together**<br>Pa: 4 4<br>Pch: 3 4 | **seprate together**<br>Ea_Pa: 3 3<br>Ea_Pch: 3 3<br>Ea_Pci: 3 3<br>Ea_Pf: 3 3 |
| Ab | **With reps**<br>Ab 2<br>IN72 5<br>Pf 6 | **With reps**<br>Ab 5<br>IN72 3<br>Sm 3<br>Pf 6 | **seprate together**<br>Fj: 3 3<br>IN72: 3 4<br>Pf: 3 3 | **seprate together**<br>Ab_IN72: 3 3<br>Pf_Ab: 2 5 |
| Pa | **With reps**<br>Ea 5<br>Pa 3<br>Pv 4 | **With reps**<br>Ea 6<br>Pa 5<br>Pv 6 | **seprate together**<br>Ea: 4 4<br>Pv: 3 5 | **seprate together**<br>Ea_Pa: 3 3<br>Pa_Pv: 3 3 |
| IN72 | **With reps**<br>Ab 5<br>IN72 3<br>Pf 4 | **With reps** | **seprate together**<br>Ab: 3 4<br>Pf: 2 2 | **seprate together**<br>Ab_IN72: 3 3 |
| Pv | **With reps**<br>Ea 5<br>Pa 4<br>Pv 3 | **With reps** | **seprate together**<br>Pa: 3 5 | **seprate together**<br>Pa_Pv: 3 3<br>Pv_Pf: 2 3 |
| Pci | **With reps**<br>Pch 2<br>Pci 2<br>IN63 5 | **With reps** | **seprate together**<br>Pch: 3 3<br>IN63: 3 4 | **seprate together**<br>Ea_Pci: 3 3<br>Pch_Pci: 2 2<br>Pci_IN63: 3 3 |
| Sm | **With reps**<br>Pch 5<br>Pf 5<br>Sm 3 | **With reps**<br>Pch 8<br>Pf 9<br>Sm 6 | **seprate together**<br>Pch: 2 2 | **seprate together**<br>Pch_Sm: 3 6 |
| IN63 | **With reps**<br>IN63 3<br>Pch 4<br>Pci 4 | **With reps** | **seprate together**<br>Pch: 3 4<br>Pci: 3 4 | **seprate together**<br>Pch_IN63: 3 3<br>Pci_IN63: 3 3 |
| Fj | **With reps**<br>Fj 2 | **With reps** | **seprate together**<br>Ab: 3 3 | **seprate together**<br>Pf_Fj: 2 2 |

***Table S3. Number of evolutionary replicates that were measured for each species in each parameter.*** *For growth and mutations, numbers indicate the number of independently evolved populations that were evolved with each partner species. For interactions and composition the comparison is between pairs that evolved together vs pairs that evolved separately (alone), and each row indicates the number of effects on a species. All strains in this table were evolved in Experiment Ev1.*

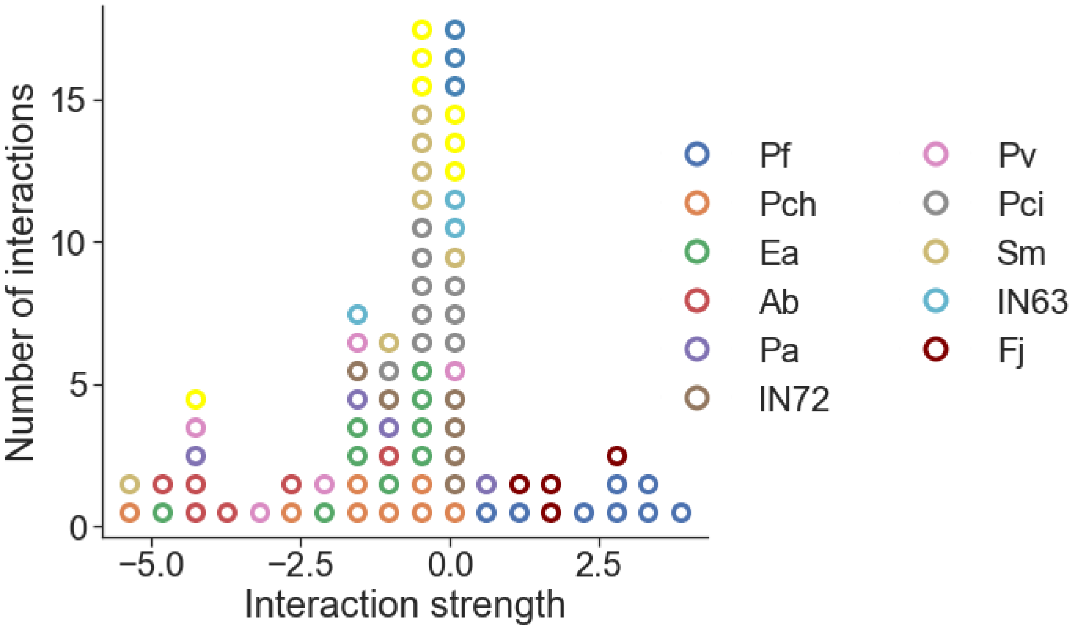


***Figure S1. Naive ancestral strains represent a wide range of interactions.*** *We quantify interactions as the effect one species has on another species' growth as the $log_2$ ratio of the abundance of a species in a specific co-culture after ~70 generations, and its abundance when grown alone for ~70 generations. Colored markers indicate the affected species, where the effects on species are binned into 20 equally-sized bins. Data from experiment Ev1.*

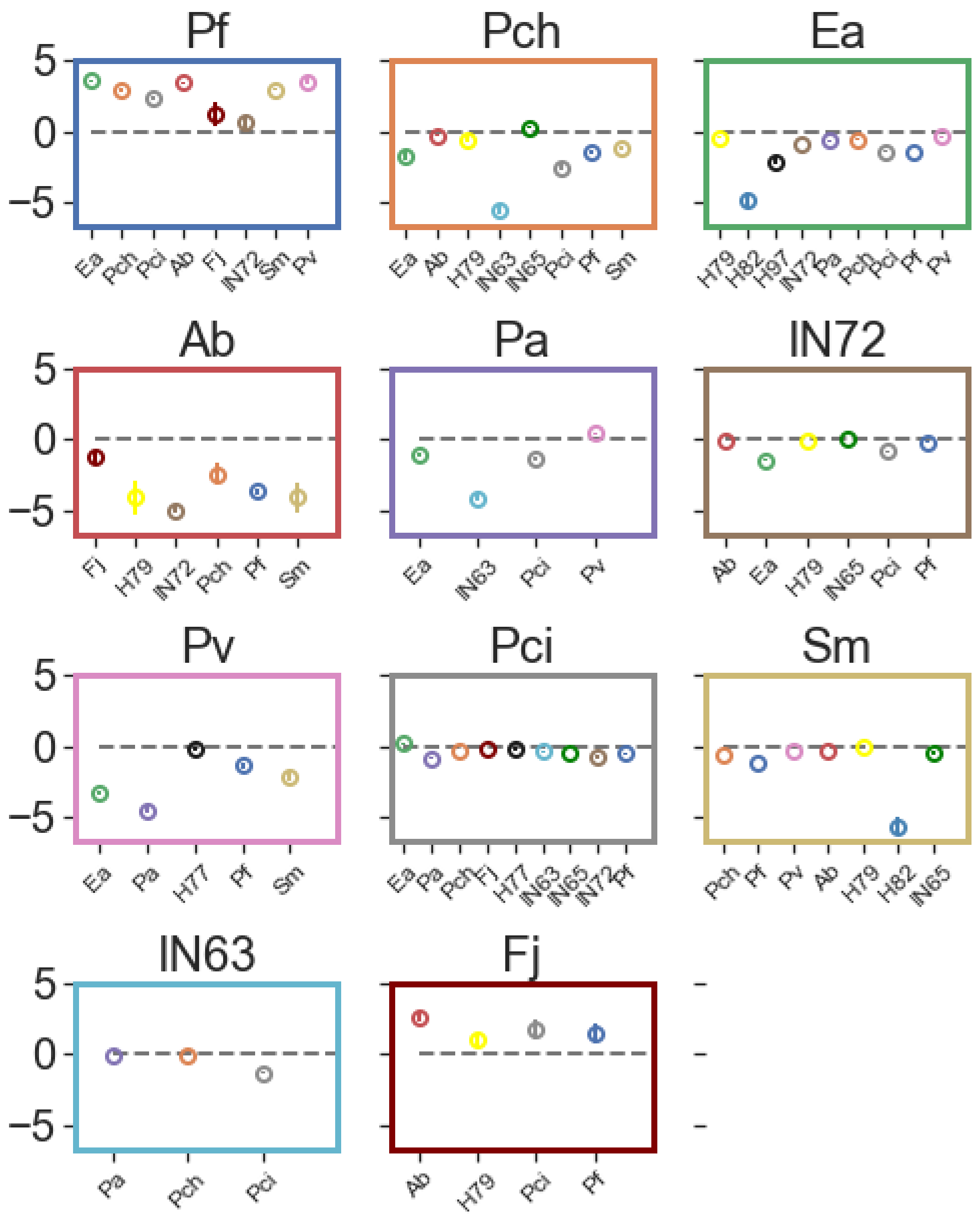


***Figure S2. Interactions of naive ancestral strains.*** *Each panel represents the effects on a specific species (title) by each of its partners (columns). Markers and error bars denote the mean and standard error of the mean of each interaction across technical replicates. Data from experiment Ev1.*

#### 1.2 Experimental setup and dataset description - pre-adapted strains

| Species | Growth<br>Number of independent populations evolved with each species | Mutations<br>Number of independent populations evolved with each species | Interactions<br>Number of independent evolved cocultures evolved in each condition | Composition<br>Number of independent evolved cocultures evolved in each condition |
|---|---|---|---|---|
| Pf | **With reps**<br>Ea 3<br>Pf 1<br>Ab 3<br>Fj 3<br>Pv 2 | **With reps**<br>Ea 5<br>Pf 2<br>Ab 5<br>Pv 4 | **seprate together**<br>Ea: 3 4<br>Ab: 3 3<br>Fj: 2 4<br>Pv: 4 4 | **seprate together**<br>Ea_Pf: 3 2<br>Pf_Ab: 2 2<br>Pf_Fj: 2 3<br>Pv_Pf: 3 2 |
| Pch | **With reps**<br>Ea 5<br>Pch 2<br>Pci 3<br>Pf 3<br>Sm 2 | **With reps**<br>Ea 4<br>Pch 3<br>Pci 3<br>Pf 4<br>Sm 5 | **seprate together**<br>Ea: 3 4<br>Pci: 3 4<br>Sm: 3 4 | **seprate together**<br>Ea_Pch: 2 5<br>Pch_IN63: 2 2<br>Pch_Pci: 2 3<br>Pch_Sm: 2 2 |
| Ea | **With reps**<br>Ea 2<br>Pa 3<br>Pch 4<br>Pci 2<br>Pf 3 | **With reps**<br>Ea 3<br>Pa 5<br>Pch 4<br>Pf 5 | **seprate together**<br>Pa: 3 4<br>Pch: 3 4<br>Pci: 3 3<br>Pf: 3 4 | **seprate together**<br>Ea_Pa: 3 3<br>Ea_Pch: 2 5<br>Ea_Pci: 3 3<br>Ea_Pf: 3 2 |
| Ab | **With reps**<br>Ab 2<br>Fj 3<br>IN72 2<br>Pf 2 | **With reps**<br>Ab 3<br>IN72 5<br>Pf 3 | **seprate together**<br>Fj: 3 4<br>IN72: 2 4<br>Pf: 3 3 | **seprate together**<br>Ab_IN72: 2 1<br>Pf_Ab: 2 2 |
| Pa | **With reps**<br>Ea 3<br>Pa 2<br>Pv 3 | **With reps**<br>Ea 5<br>Pa 3<br>Pv 5 | **seprate together**<br>Ea: 3 4<br>Pv: 4 4 | **seprate together**<br>Ea_Pa: 3 3<br>Pa_Pv: 3 3 |
| IN72 | **With reps**<br>Ab 2<br>Fj 1<br>IN72 2<br>Pf 2 | **With reps** | **seprate together**<br>Ab: 3 4 | **seprate together**<br>Ab_IN72: 2 1 |
| Pv | **With reps**<br>Pa 3<br>Pv 2<br>Pf 2 | **With reps** | **seprate together**<br>Pa: 4 4<br>Pf: 4 4 | **seprate together**<br>Pa_Pv: 3 3<br>Pv_Pf: 3 2 |
| Pci | **With reps**<br>Ea 3<br>Pch 3<br>Pci 2<br>IN63 1<br>Pf 1 | **With reps** | **seprate together**<br>Ea: 3 3<br>Pch: 3 4<br>IN63: 3 2 | **seprate together**<br>Ea_Pci: 3 3<br>Pch_Pci: 2 3<br>Pci_IN63: 3 3 |
| Sm | **With reps**<br>Pch 2<br>Sm 2 | **With reps** | **seprate together**<br>Pch: 3 4 | **seprate together**<br>Pch_Sm: 2 2 |
| IN63 | **With reps**<br>IN63 3<br>Pch 1<br>Pci 1 | **With reps** | **seprate together**<br>Pci: 3 2 | **seprate together**<br>Pch_IN63: 2 2<br>Pci_IN63: 3 3 |
| Fj | **With reps**<br>Ab 3<br>Fj 2<br>Pf 3 | **With reps** | **seprate together**<br>Ab: 3 4<br>Pf: 2 4 | **seprate together**<br>Pf_Fj: 2 3 |

***Table S4. Number of evolutionary replicates that were measured for each species in each parameter.*** *For growth and mutations, numbers indicate the number of independently evolved* *populations that were evolved with each species. For interactions and composition the comparison is between pairs that evolved together vs pairs that evolved separately (alone), and each row indicates the number of effects of a species. All strains in this table were evolved in Experiment Ev2.*

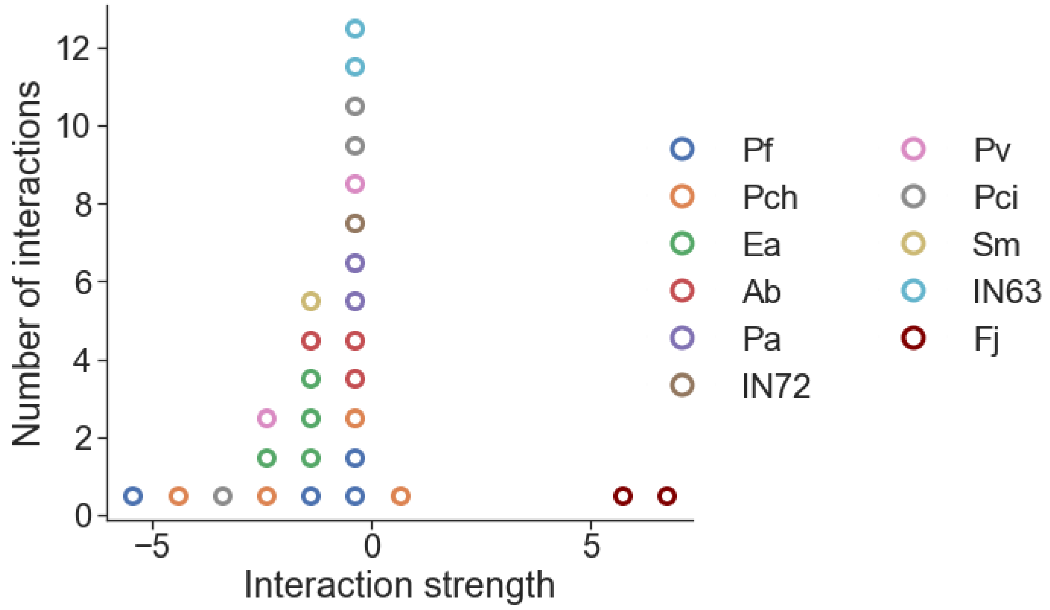


***Figure S3. Distribution of interactions of pre-adapted ancestral strains.*** *We quantify interactions as the effect one species has on another species growth as the $log_2$ ratio of the abundance of a species in a specific co-culture after ~70 generations, and its abundance when grown alone for ~70 generations. Colored markers indicate the affected species. Data from experiment Ev2.*

#### 2. Supplementary data for naive strains and pairs

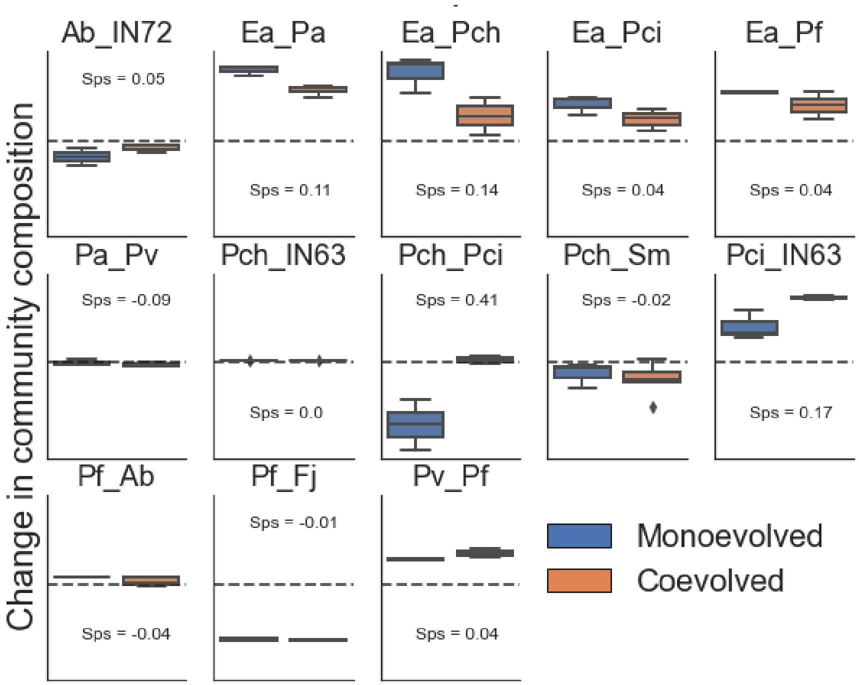


***Figure S4. Change in community composition of naive pairs.*** *Each panel represents the change in the community composition of a pair when the strains composing it were evolved separately as monocultures (Blue) or coevolved in co-culture (orange). Change in composition is measured as the fraction of a species in the evolved co-culture minus its fraction in the ancestral co-culture. Boxes indicate the quartiles and whiskers are expanded to include values no further than 1.5X interquartile range of independently evolved co-cultures. Sps indicates each pair's specificity score. Data from experiment Ec1.*

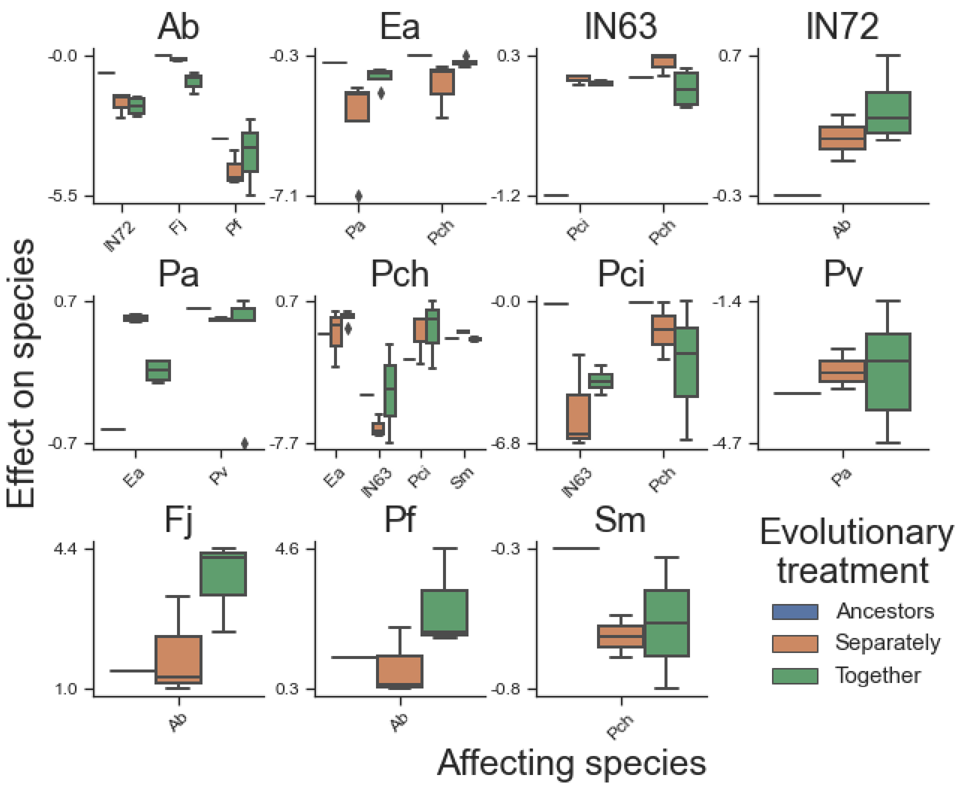


***Figure S5. Interaction between strains evolved from naive ancestor.*** *Each panel represents the effects of various species on the species indicated in the title. Grouping on the x-axis are the different affecting species, and box color represents different evolutionary treatments. Boxes indicate the quartiles of independently evolved co-cultures and whiskers are expanded to include values no further than 1.5X interquartile range. Data from experiments Ec2-5.*

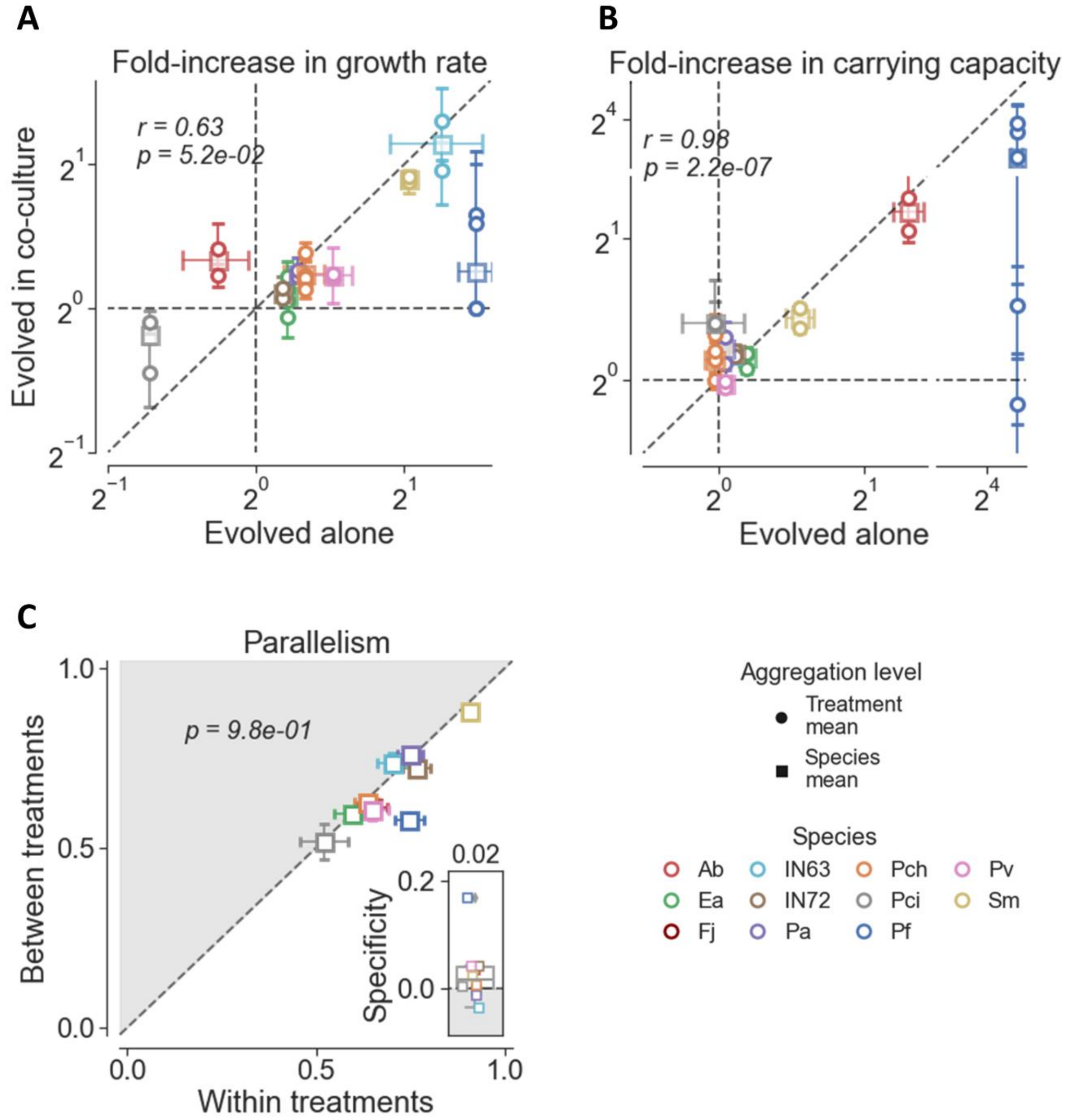


**Figure S6: Evolution of growth abilities** of naive strains**.** (A, B) Fold-increase in growth rate (A) and fold increase in productivity (B) of strains that evolved alone vs strains that evolved in coculture. Colors indicate different species, squares denote the mean of all coevolved strains of the same species (Species grand mean), and circles denote all coevolved strains of the same species that evolved with the same partner (treatment mean). Error bars indicate the standard error of the mean of 2-5 independently evolved strains that evolved in a specific treatment (Table S4). (C) Parallelism in growth parameters of strains that evolved in the treatment against that of strains that evolved in different treatments. Squares denote the mean of each species across treatments and error bars indicate the standard error of the mean. Inset shows the distribution specificity scores in the evolution of growth parameters, and the number above is the median score across all species. Fisher's method for combining p-values for the null hypothesis tha distances within treatments are not smaller: p-value = 0.007 . Data from experiment Gr1. Statistics in panels A and B are Pearson r and associated p, statistic in panel C is a one-sided Wilcoxon test p-value.

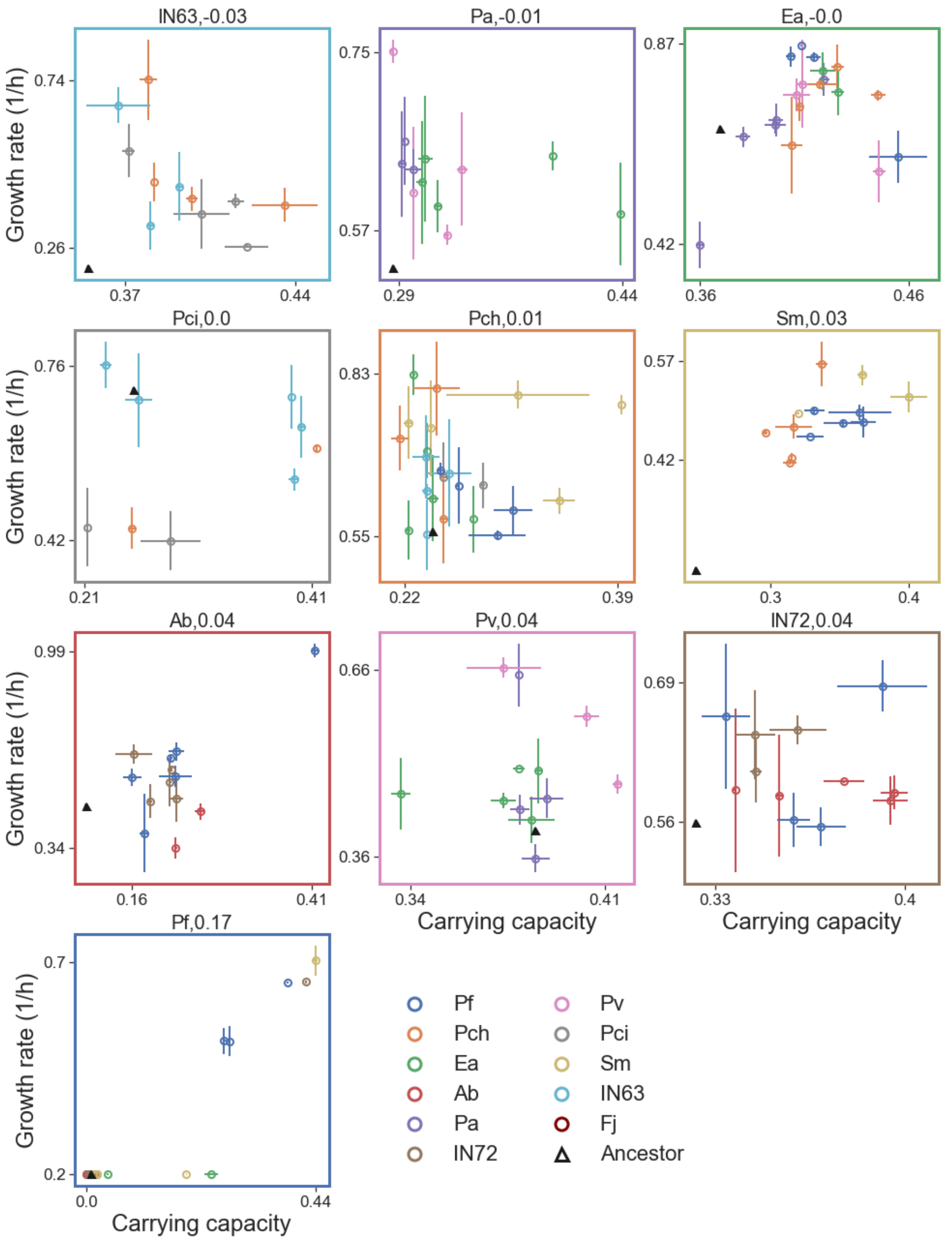


***Figure S7: Evolution of growth abilities of naive strains.*** *Plots of the carrying capacity against the growth rate of each species after evolving in multiple biotic conditions. Triangles represent the ancestral strains, and each circle corresponds to an evolved strain. Colors represent the partner with which species' evolved. Circles and error bars are means and standard errors of 2 technical replicates. Numbers in the titles are the parallelism score of each species. Data from experiment Gr1.*

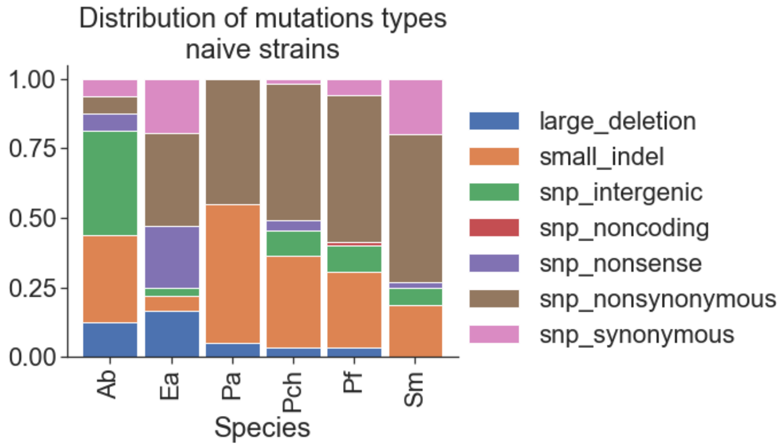


***Figure S8: Distribution of mutation types in naive strains.*** *Each bar represents the full distribution of mutations in all the strains of a species that were evolved across all treatments.*

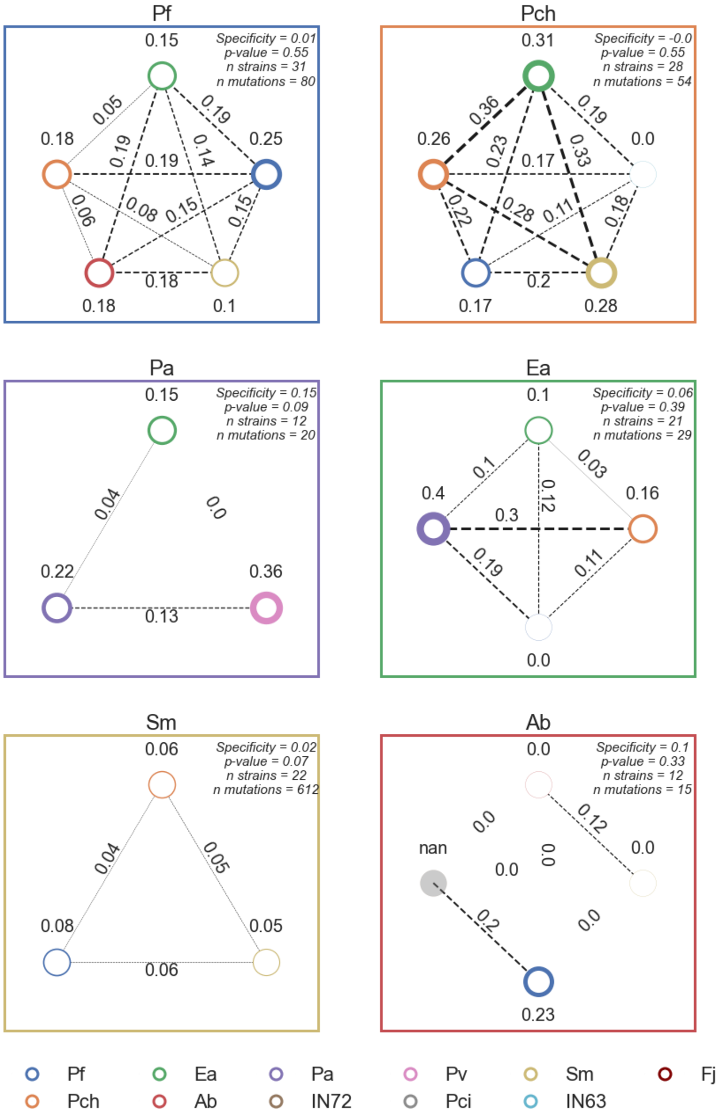


***Figure S9: Pairwise comparisons between the parallelism in mutated genes within treatments and the parallelism between treatments.*** *Each panel summarizes all the comparisons of gene-level parallelism between strains of a species (panel title). Each circle represents a treatment (biotic partner, legend colors), and the number above indicates the mean parallelism within this treatment. Connecting lines and the numbers above them indicate the mean parallelism between each pair of treatments. Circle and line thickness indicate the mean parallelism within and between treatments respectively. Numbers in the right upper corner give the specificity score (parallelism within - parallelism between); p-value generated by a permutations test for the null hypothesis that there is no difference between the parallelism within and between treatments and after Bonferroni correction for multiple comparisons; number of overall strains tested in these comparison, this excludes strains that had no mutations or only synonymous mutations; and the number of mutations that generated these comparisons.*

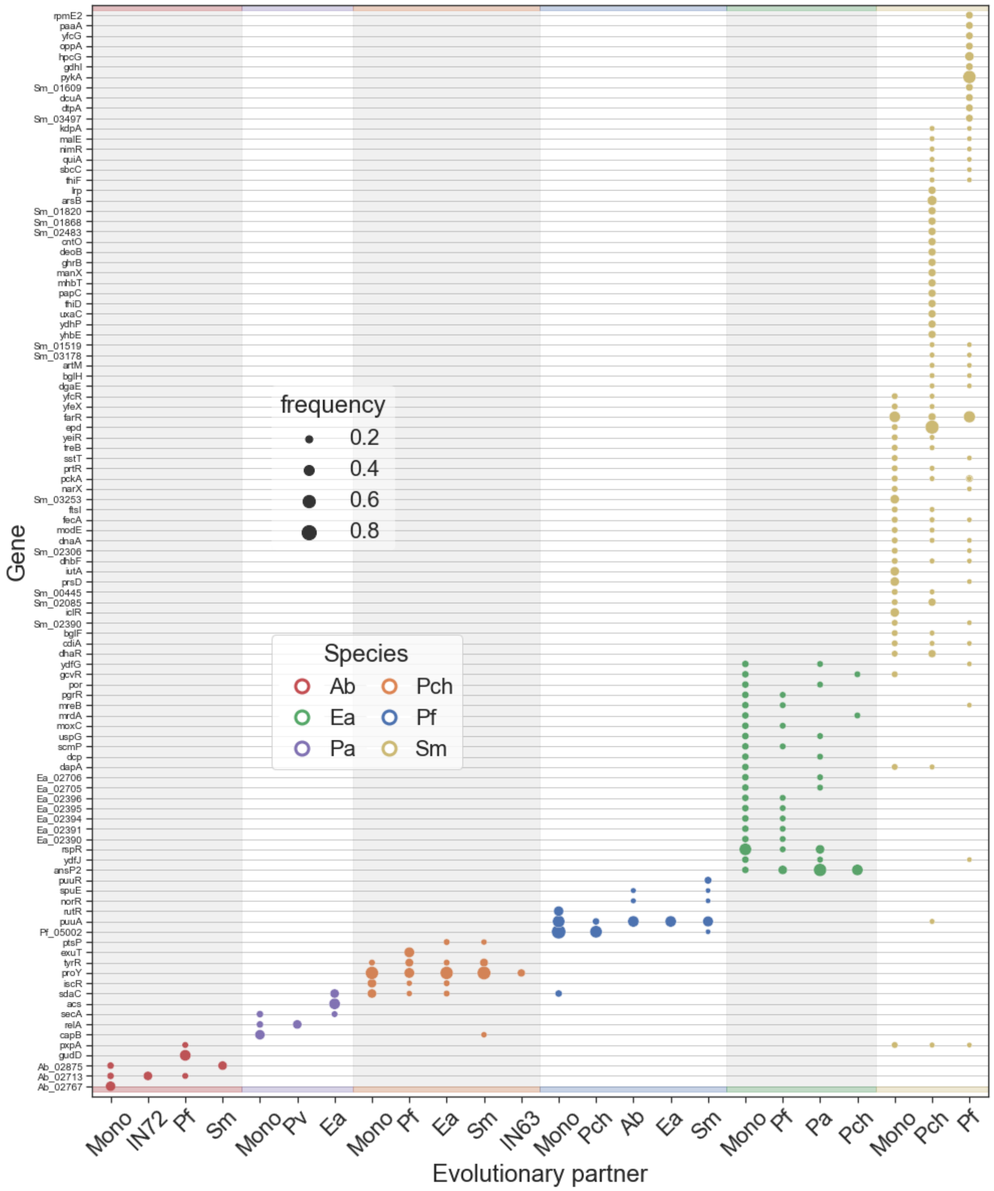


***Figure S10: Parallelly mutated genes in naive strains for each species**. Genes in this plot are all genes that were mutated in more than one strain. Each color is a different species and columns indicate the biotic context it evolved in. Marker size indicates the fraction of strains that evolved in the specific biotic context that were mutated in this gene. Genes written in the form "species_number" are hypothetical proteins. By this criterion, 61% of the genes that were mutated in more than one strain are mutated in more than one treatment.*

| Species | Partner (treatment) | Affected gene | Number of strains mutated within treatment | Number of strains within treatment | Number of strains mutated in other treatment | Total number of strains in other treatments | Boschloo test p-value | Gene product |
|---|---|---|---|---|---|---|---|---|
| Ab | Pf | gudD | 3 | 6 | 0 | 11 | 0.021811 | Putative glucarate transporter/Glucarate dehydratase |
| Pa | Ea | acs_2 | 3 | 6 | 0 | 11 | 0.021811 | Acetyl-coenzyme A synthetase |
| Pch | Pch | exuT_3 | 3 | 7 | 0 | 23 | 0.006047 | Hexuronate transporter |
| Pf | Pf | rutR_2 | 2 | 5 | 0 | 28 | 0.013981 | HTH-type transcriptional regulator RutR |
| Pf | Pf | Pf_05002 | 4 | 5 | 4 | 28 | 0.005483 | Hypothetical protein |
| Sm | Pch | epd_2 | 6 | 8 | 1 | 15 | 0.001239 | D-erythrose-4-phosphate dehydrogenase |
| Sm | Pf | pykA | 6 | 9 | 0 | 14 | 0.000409 | Pyruvate kinase II |

***Table S5: Extended table of genes that were mutated predominantly in one evolutionary treatment.*** *Table includes all mutated genes in naive strains that had a p-value < 0.05 in a Boschloo test before corrections for multiple comparisons.*

#### 3. Supplementary data for pre-adapted strains and pairs

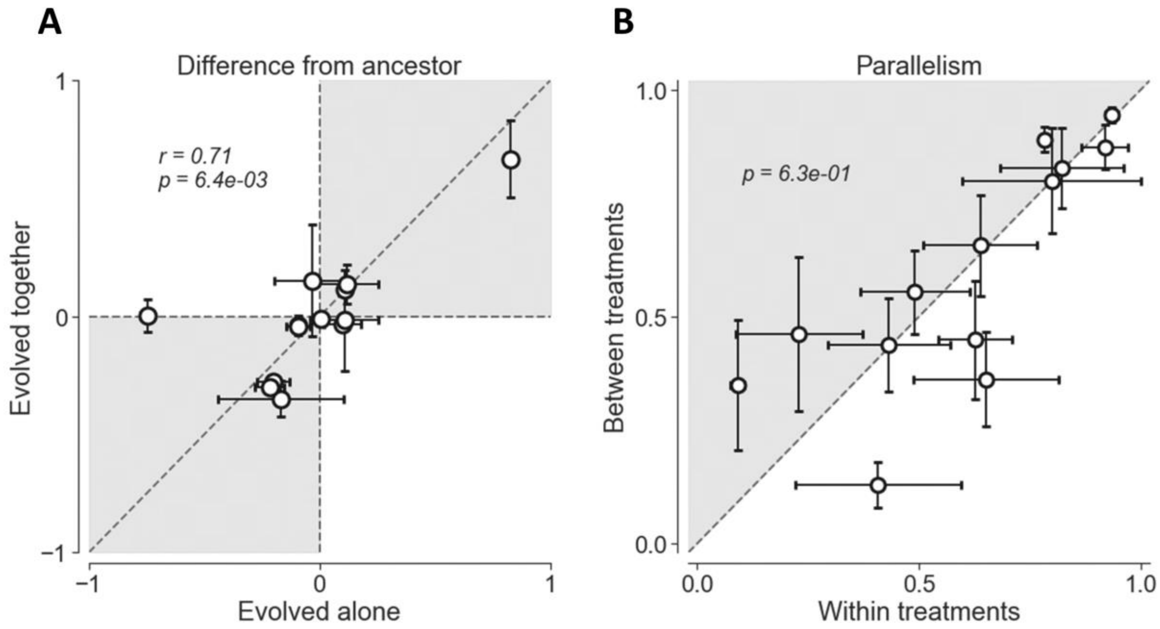


***Figure S11: Evolution of composition in pre-adapted pairs.*** *Change in the composition of pairs of strains that evolved together, against the composition after they evolved separately in monocultures. Change in composition is measured as the fraction of a species in the evolved co-culture minus its fraction in the ancestral co-culture. Each circle represents a unique and initially identical pair of species, and the circle center and error bars represent the mean and the standard error of 2-6 independently evolved co-cultures. (B) Parallelism in the evolution of composition within treatment (parallelism between pairs of strains that either evolved alone or coevolved) against the parallelism between treatments (parallelism between pairs strains that evolved alone to strains that coevolved). Circles and error bars indicate the mean and the standard error of each unique pair of species. Data from experiment Ec1. Statistics in panels A are Pearson r and associated p, statistic in panel B is a one-sided Wilcoxon test p-value.*

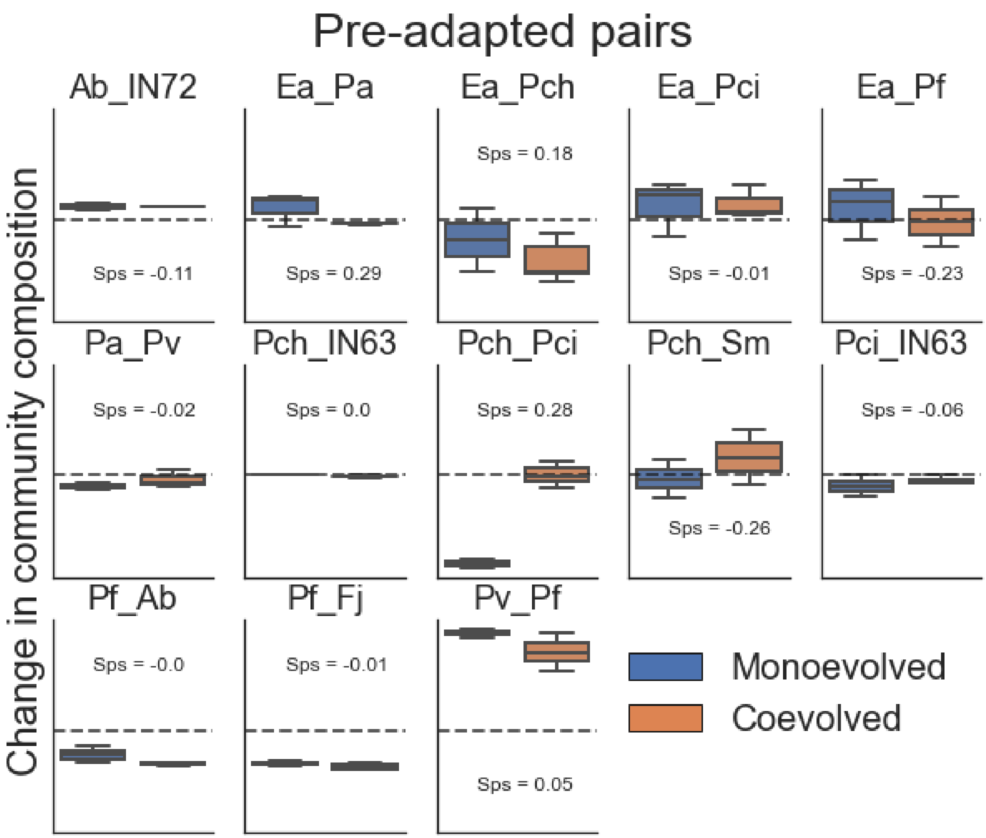


***Figure S12. Change in community composition of pre-adapted pairs.*** *Each panel represents the change in the community composition of a pair when the strains composing it were evolved separately as monocultures (Blue) or coevolved in co-culture (orange). Change in composition is measured as the fraction of a species in the evolved co-culture minus its fraction in the ancestral co-culture. Boxes indicate the quartiles and whiskers are expanded to include values no further than 1.5X interquartile range of independently evolved co-cultures. Sps indicates each pair's specificity score. Data from experiment Ec1.*

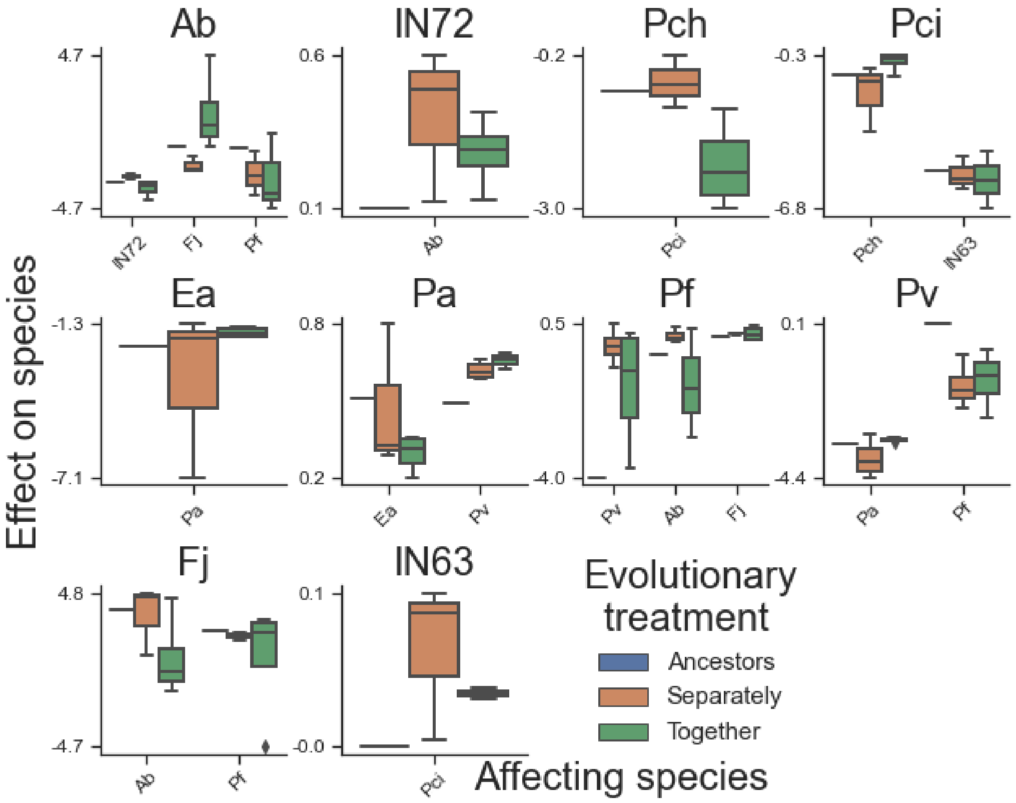


***Figure S13. Interactions between pre-adapted strains.*** *Each panel represents the effects of various species on the species indicated in the title. Grouping on the x-axis are the different affecting species, and box color represents different evolutionary treatments. Boxes indicate the quartiles of independently evolved co-cultures and whiskers are expanded to include values no further than 1.5X interquartile range. Data from experiments Ec2-5.*

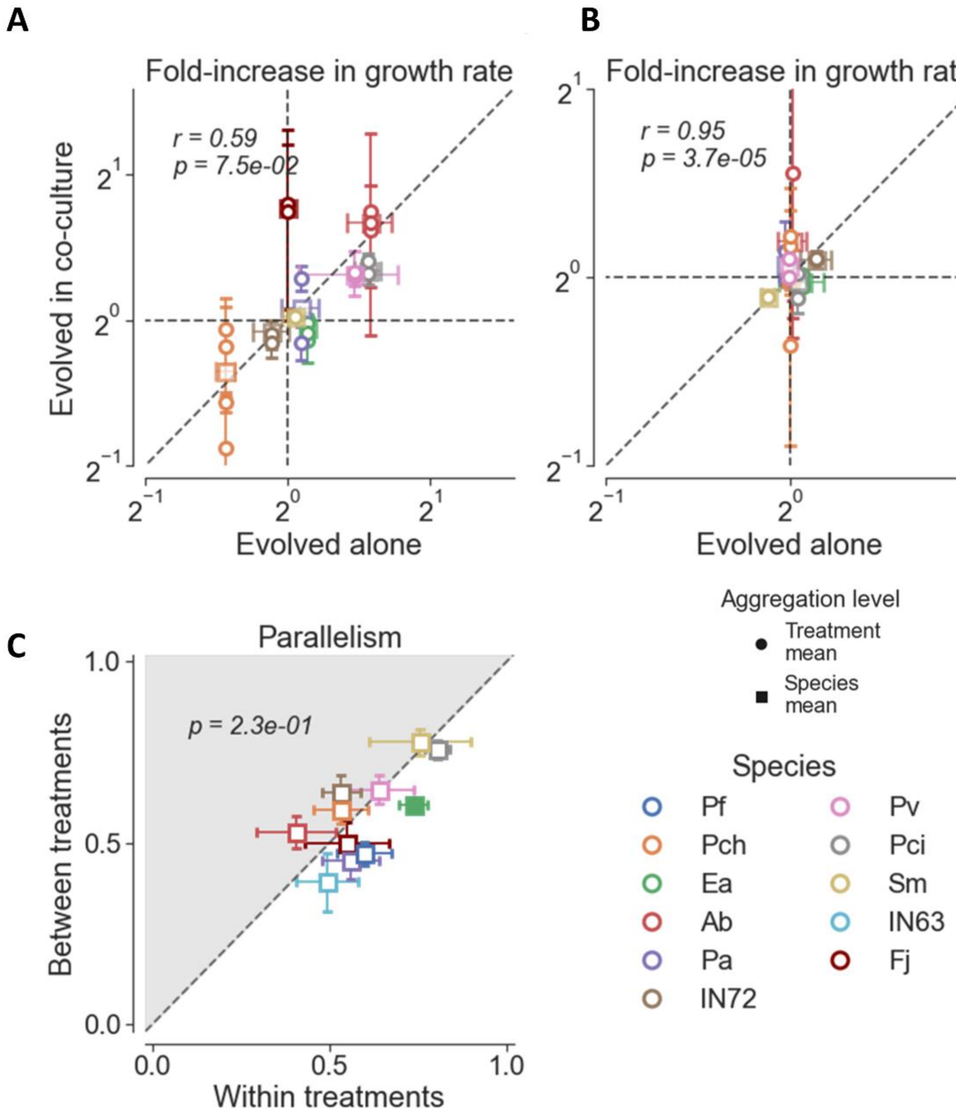


***Figure S14: Evolution of growth abilities of pre-adapted strains.*** *(A, B) Fold-increase in growth rate (A) and fold increase in productivity (B) of strains that evolved alone vs strains that evolved in coculture. Colors indicate different species, squares denote the mean of all coevolved strains of the same species (Species grand mean), and circles denote all coevolved strains of the same species that evolved with the same partner (treatment mean). Error bars indicate the standard error of the mean of independently evolved strains that evolved in a specific treatment. (C) Parallelism in growth parameters of strains that evolved in the treatment against that of strains that evolved in treatments. Squares denote the mean of each species across treatments and error bars indicate the standard error of the mean. Fully colored squares indicate that a species had significantly higher parallelism within treatments in a permutations test after correcting for multiple Bonferroni correction for multiple comparisons with a false discovery rate of 5%. Fisher's method for combining p-values for the null hypothesis tha distances within treatments are not smaller: p-value = 0.007 . Data from experiment Gr1. Statistics in panels A and B are Pearson r and associated p-value, statistic in panel B is a one-sided Wilcoxon test p-value.*

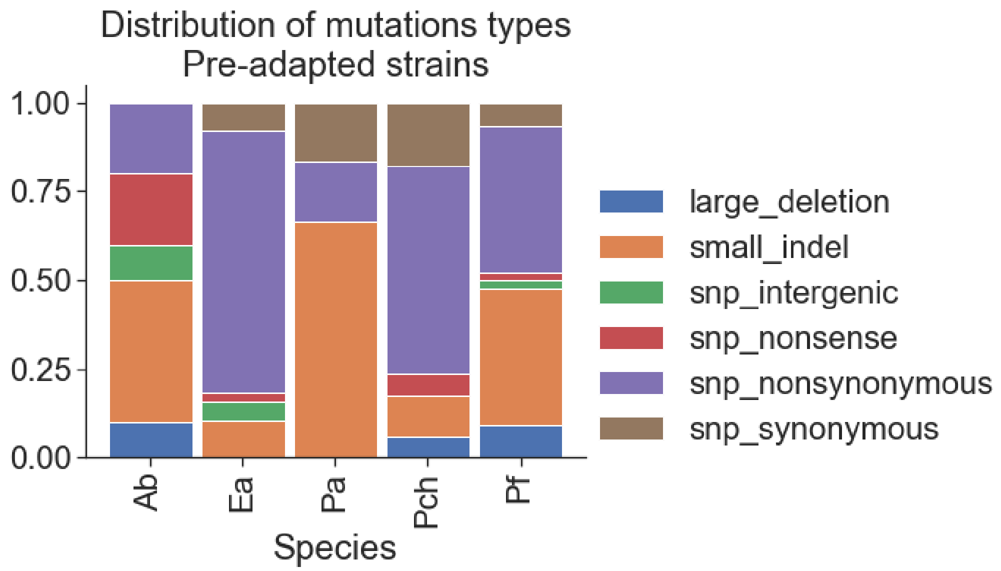


***Figure S15: Distribution of mutation types of Pre-adapted strains.*** *Each bar represents the full distribution of mutations a species had across all treatments.*

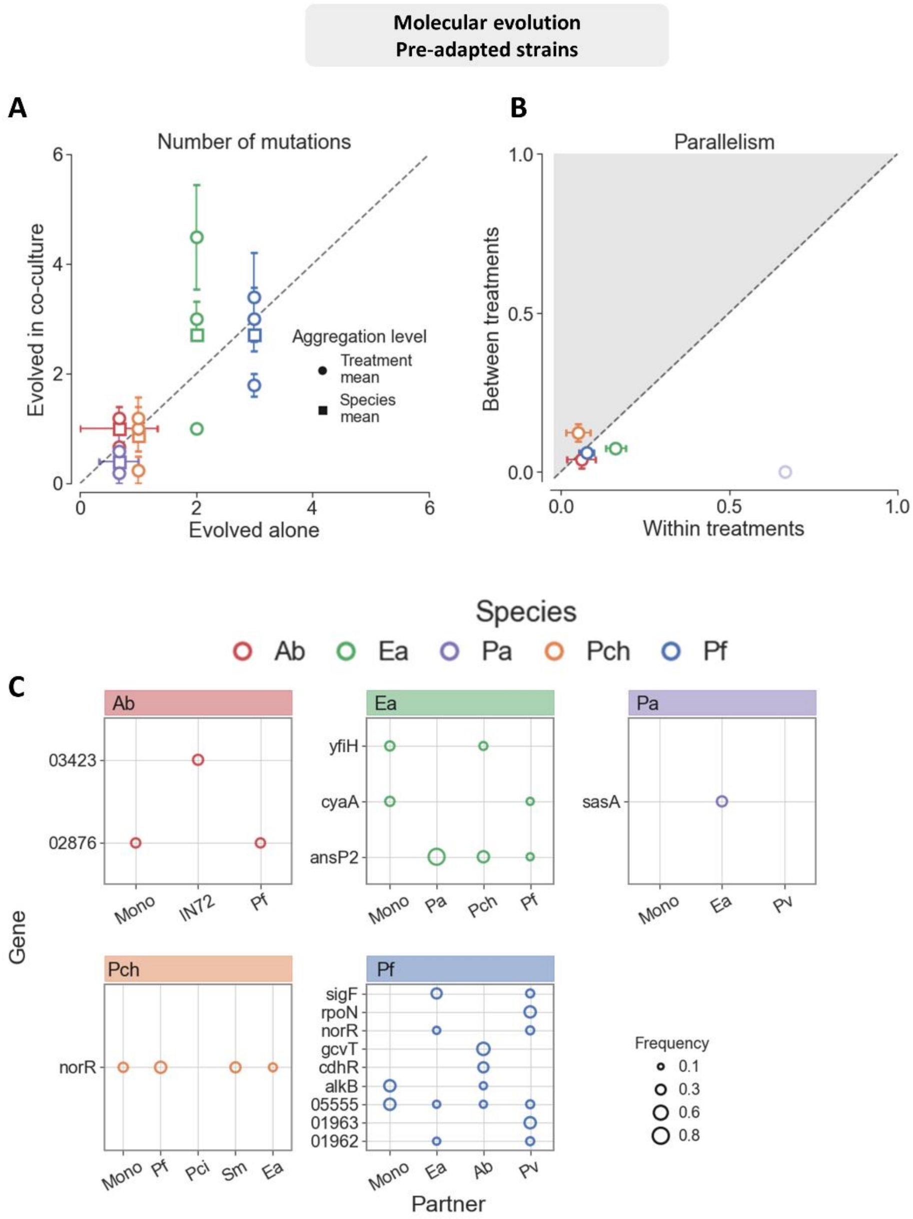


***Figure S16. Genome evolution in pre-adapted strains*** *(A) Mean number of mutations per strain of species when evolved alone vs when evolved in co-culture. Squares indicate the mean number across all strains of the same species that evolved in co-culture (grand mean), and circles indicate strains that evolved with a specific partner (treatment mean). Error bars denote the standard error mean. Wilcoxon p-value = 0.81. (B) Parallelism between mutated genes in strains evolved in the same treatment, against the Parallelism between strains that evolved in treatments. Parallelism in genomic evolution is measured as the dice similarity in mutated genes. Error bars indicate the standard error mean for each species. P-value after combining p-values using Fisher's method = 0.14 . (C) Parallelly mutated genes in each species. Genes in this plot were mutated in more than one strains. Each color is a different species and columns indicate the biotic context it evolved in. Marker size indicates the fraction of strains that evolved in the specific biotic context*

*that were mutated in this gene. Genes names written as numbers are hypothetical proteins.*

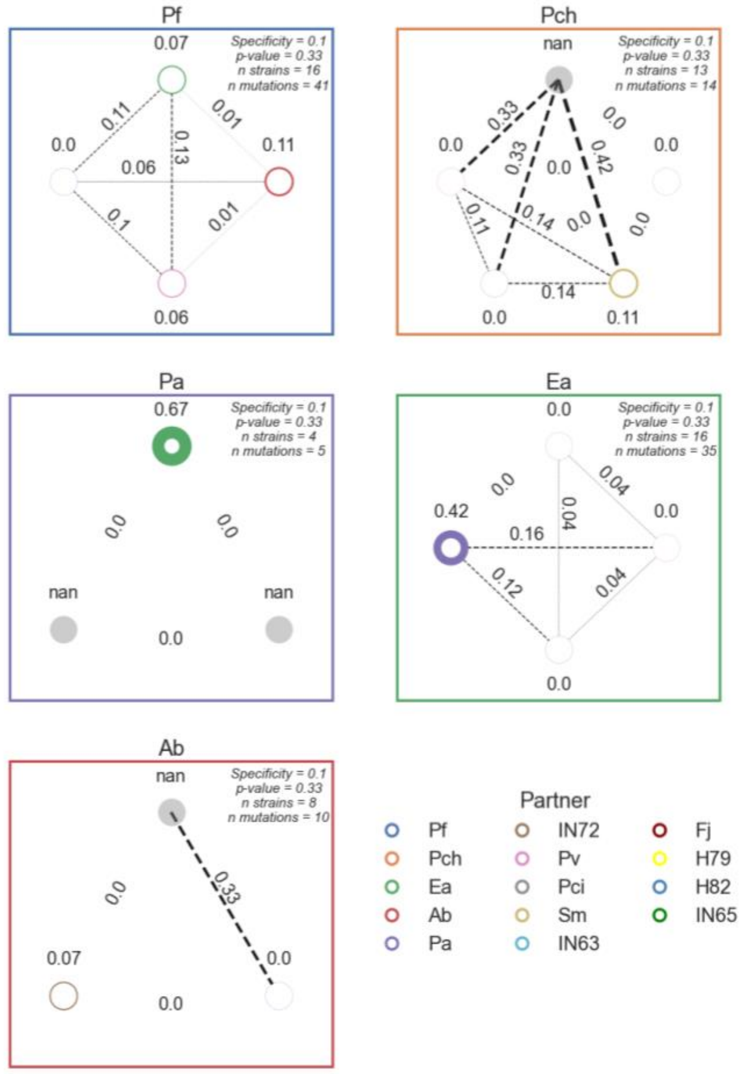


***Figure S17: Pairwise comparisons between the parallelism in mutated genes within treatments and the parallelism between treatments in pre-adapted strains.*** *Each panel summarizes all the comparisons of gene-level parallelism between strains of a species (panel title). Each circle represents a treatment (biotic partner, legend colors), and the number above indicates the mean parallelism within this treatment. Connecting lines and the numbers above them indicate the mean parallelism between each pair of treatments. Circle and line thickness indicate the mean parallelism within and between treatments respectively. Numbers in the right upper corner give the specificity score (parallelism within - parallelism between); p-value generated by a permutations test for the null hypothesis that there is no difference between the parallelism within and between treatments and after Bonferroni correction for multiple comparisons; number of overall strains tested in these comparison, this excludes strains that had no mutations or only synonymous mutations; and the number of mutations that generated these comparisons.*

| Species | Evolved community | Affected gene | Number of strains mutated within treatment | Number of strains within treatment | Number of strains mutated in other treatment | Total number of strains in other treatments | Boschloo test p-value | Gene product |
|---|---|---|---|---|---|---|---|---|
| *Ea* | *Ea_Pa* | *ansP2* | *5* | *5* | *3* | *12* | *0.005341* | *L-asparagine permease 2* |
| *Pf* | *Pf_Ab* | *gcvT_2* | *3* | *5* | *0* | *11* | *0.010982* | *Aminomethyltransferase* |
| *Pf* | *Pv_Pf* | *Pf_01963* | *2* | *4* | *0* | *12* | *0.036403* | *Hypothetical* |
| *Pf* | *Pv_Pf* | *rpoN* | *2* | *4* | *0* | *12* | *0.036403* | *RNA polymerase sigma-54 factor* |

***Table S6: Extended table of genes that were mutated predominantly in one evolutionary treatment.*** *Table includes all mutated genes in pre-adapted strains that had a p-value<0.05 in a Boschloo test before corrections for multiple comparisons.*

#### 4. Comparison between naive and pre-adapted strains

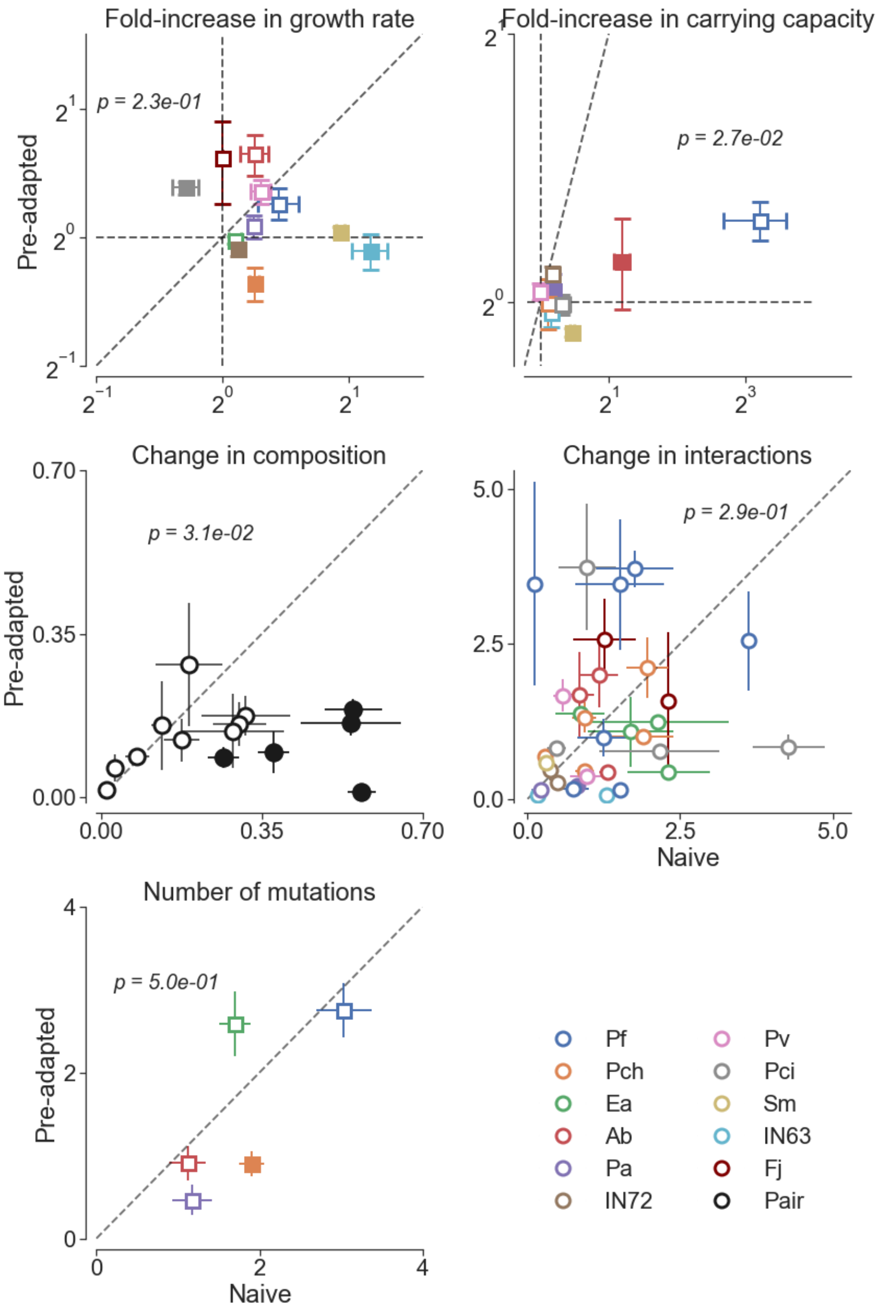


***Figure S18: The rate of adaptation decreases in pre-adapted strains.*** *Each panel shows the mean and standard error of change in naive strains against pre-adapted strains. Filled markers signify species that changed significantly more when evolved in one of the treatments (Mann-Whitney U-test after Bonferroni correction for multiple comparisons with a 5% FDR). Text on the plots indicates the one-sided Wilcoxon test p-value with the hypothesis that naive strains or pairs changed more than pre-adapted strains or pairs.*

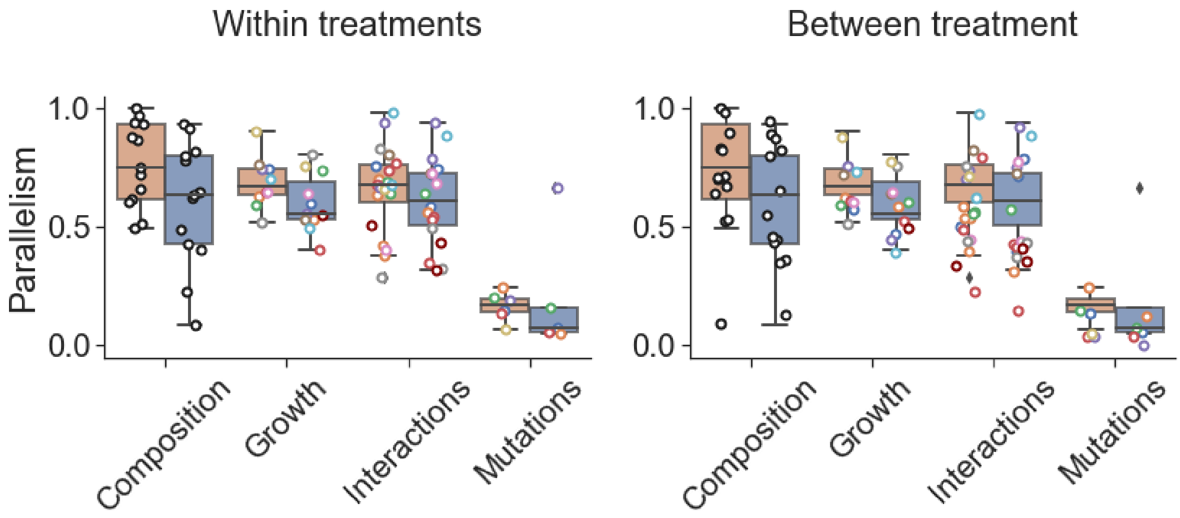


***Figure S19: Parallelism is lower in pre-adapted strains****. Parallelism within treatment (left) and between treatments (right) in naive (orange) and pre-adapted (blue) strains. One-sided Wilcoxon test p-value = 0.008. Dots indicate the mean for each species (colored) or co-culture (black).*

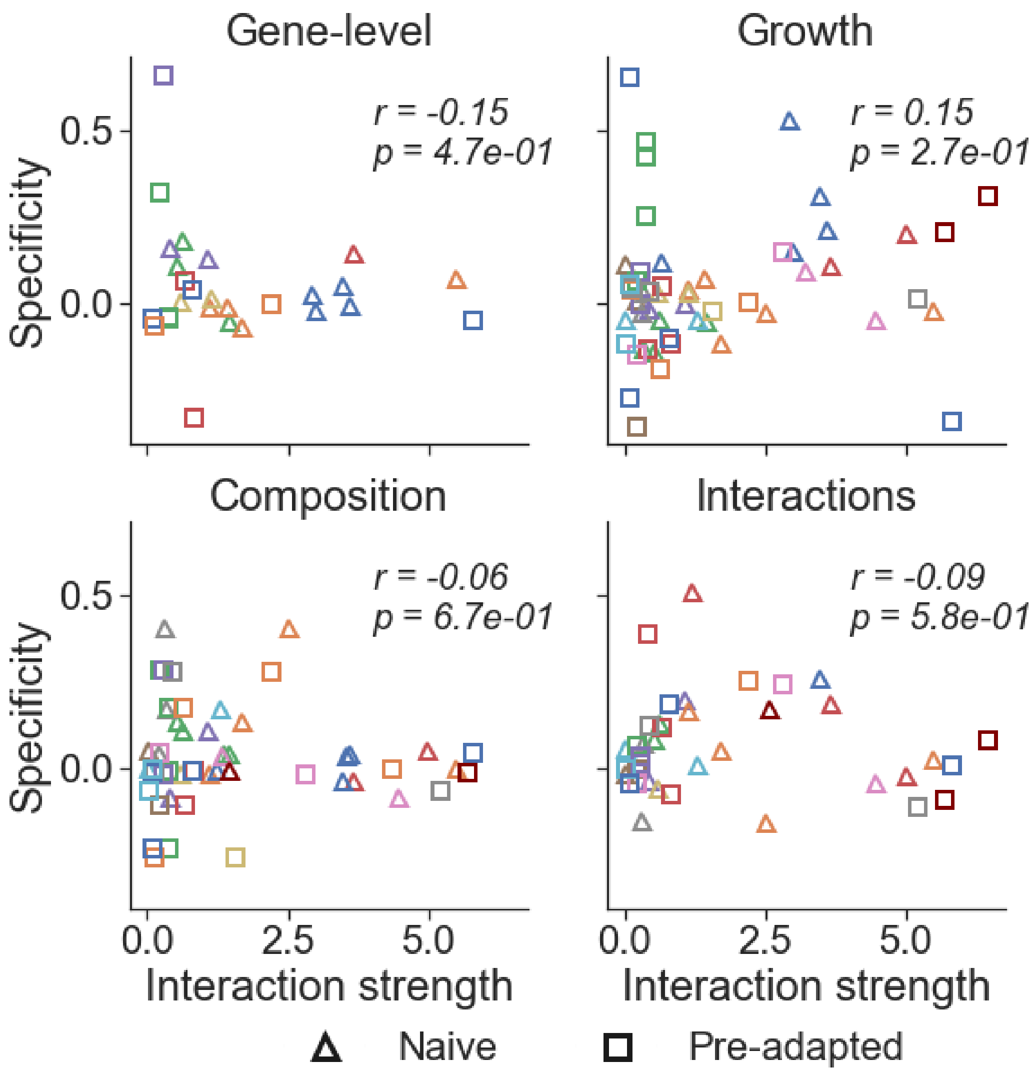


***Figure S20: Specificity does not correlate with interaction strength**. Specificity is calculated for each species between its strains that evolved alone and strains that evolved with each different partner. This is plotted against the effect (one-way interaction) of the partner on the species.. Triangles are naive evolved species (evolved in Ev1), and squares are pre-adapted evolved species (evolved in Ev2). Each panel represents the specificity in the evolution of a different trait. Text on the plots indicates the r, and p, of* Pearson *correlation.*

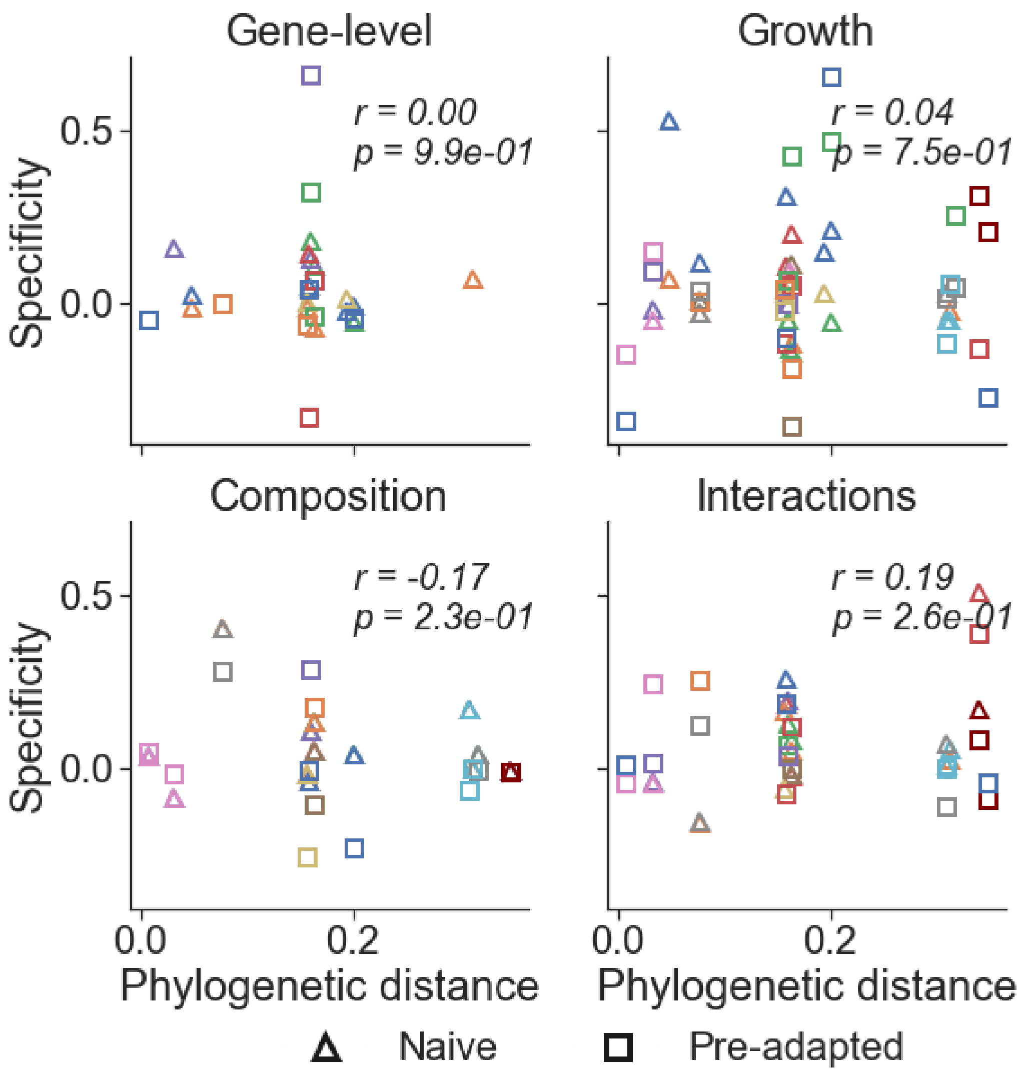


***Figure S21: Specificity does not correlate with Phylogenetic distance from partner***. *Specificity is calculated for each species between its strains that evolved alone to strains that evolved with each different partner. This is plotted against the phylogenetic distance between the species. Phylogenetic distance is calculated as the per-site substitution rate in species 16S rRNA sequence. Triangles are naive evolved species (evolved in Ev1), and squares are pre-adapted evolved species (evolved in Ev2). Each panel represents the specificity in the evolution of a different trait. Text on the plots indicates the r and p of Pearson correlation.*

#### 5. Parallelism quantification

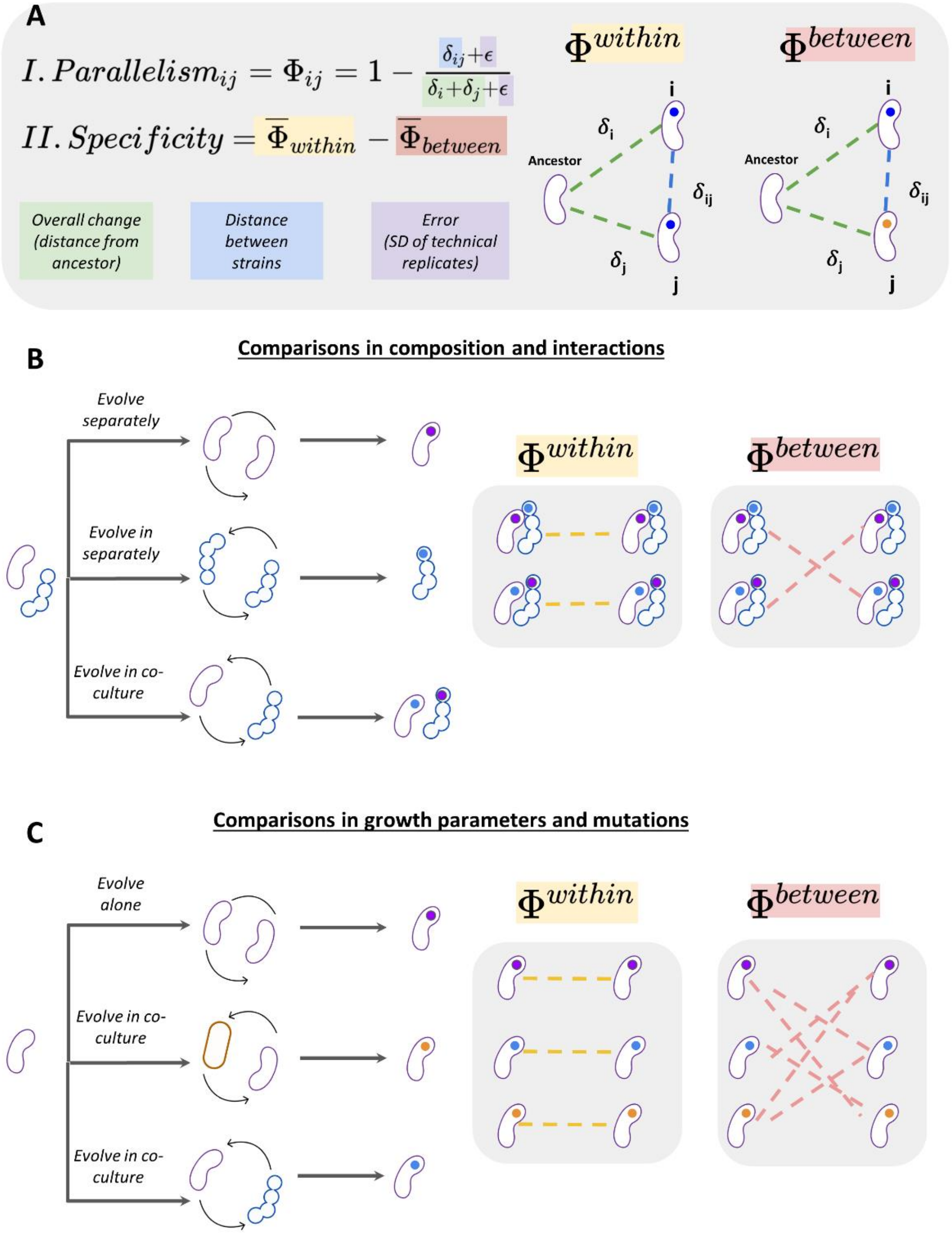


***Figure S22. Measures and comparisons used throughout the study.*** *(A) Parallelism is defined as the fraction of the overall change which is similar between two strains. Specificity is defined as the mean parallelism of strains, or pairs of strains, that evolved independently in the same treatment, minus the parallelism strains that evolved independently in different treatments. (B, C) We compare the parallelism within treatment to the parallelism between treatments. (B) When we measure co-culture properties (composition and interactions), the comparison is between*

*pairs of strains that co-evolved together, to pairs of strains that were evolved separately. (C) Comparison in growth parameters and mutations is done between multiple treatments, where each treatment is a specific community context - monoculture, co-culture A, co-culture B, etc.*

We devised a measure of parallelism which quantifies how much of the total amount of change in each trait, or set of traits, is shared between two strains. This measure is used rather than simple distances, in order to make the quantitative results easier to interpret, and because it is equivalent to the Dice similarity coefficient which is standardly used for quantifying parallelism at the genomic level. In the following section, we discuss how this measure was devised and its behavior in different conditions. Finally, we also show that our main findings would not have changed qualitatively if simple distances were used.

For each pair of strains $i$ and $j$ that share a common ancestor, we define $\delta_{ij}$ as the distance between them, and $\delta_i$, $\delta_j$ as the distance of $i, j$ from the shared ancestor. Therefore, for every $i$, $j$, $\delta_{ij} \leq (\delta_i + \delta_j)$, and $(\delta_i + \delta_j)$ could be defined as the total amount of change relative to the ancestor. Thus, Parallelism $(\Phi_{ij})$, the fraction of the total amount of change that is shared between a pair of strains, is quantified as $\Phi_{ij} = 1 - \frac{\delta_{ij}}{(\delta_i+\delta_j)}$

In this case, $0 \leq \Phi_{ij} \leq 1$. When $\delta_{ij} = 0$ , strains share all of the change and $\Phi_{ij} = 1$, and when two strains share non of the change $\Phi_{ij} = 0$.

$\Phi_{ij} = 0$ occurs when two strains evolved in a trait, or a set of traits, to exactly opposing directions. While it rarely occurs in our dataset, this might produce some non-intuitive results when the total amount of change $(\delta_i + \delta_j)$ is small, and differences between the strains could be attributed to measurement or biological noise. To diminish this bias, modified the calculation of $\Phi_{ij}$ to account for measurement noise $(\epsilon)$, which is calculated as the standard error of the mean between technical replicates in each trait, such that $\Phi_{ij} = 1 - \frac{\delta_{ij}+\epsilon}{(\delta_i+\delta_j)+\epsilon}$.

As long as $(\delta_i + \delta_j) \gg \epsilon$, the total amount of change can not be attributed to noise, and $\Phi_{ij}$ is not significantly affected by the addition of $\epsilon$. However, as $(\delta_i + \delta_j) \to \epsilon$, $\Phi_{ij} \to 1$, such that two strains that remain similar to their common ancestor are assigned $\Phi_{ij} \approx 1$, if indistinguishable.

The Dice similarity coefficient is equivalent to our measure of parallelism, when $\delta_i, \delta_j, \delta_{ij}$ are measured using Hamming distances. Dice similarity is commonly defined as $\frac{2|G_i \cap G_j|}{|G_i| + |G_j|}$, where $G_i$, and $G_j$ are the genes mutated in strain $i$, and $j$ respectively. By definition, the genes are mutated with respect to their ancestor and therefore $(\delta_i + \delta_j) = |G_i| + |G_j|$, and $\delta_{ij} = |G_i| + |G_j| - 2|G_i \cap G_j|$ .

Below, we show that the results are not qualitatively different if distances, rather than parallelism, are used. First, we compare the distances between treatments and within treatments, as done throughout the paper with parallelism scores. A similar trend emerges - in each trait the distance between treatments are typically slightly larger and typically remain close to the 1:1 line (Figure S23). Next, we compare the specificity scores calculated when using parallelism ($\underline{\Phi_{within}} - \underline{\Phi_{between}}$) to specificity scores calculated when using distances ($\underline{\delta_{between}} - \underline{\delta_{within}}$). While specificity scores calculated in terms of distances are less interpretable quantitatively and their scale varies between traits, these correlate well with their parallelism based equivalents (Figure S24).

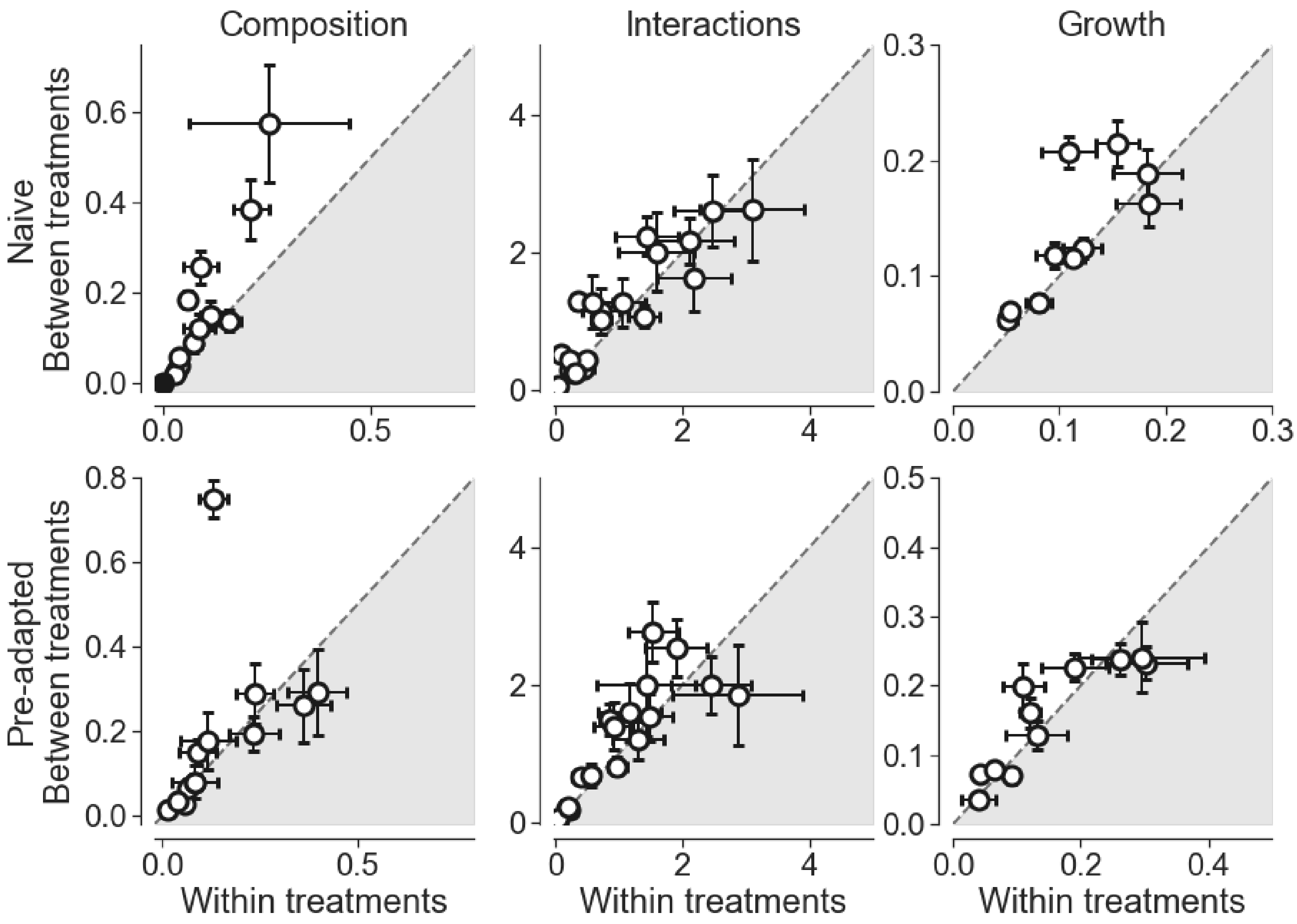


***Figure S23. Evolution of coculture properties is similar regardless of coevolution.*** *Difference in composition (A) and interactions (B) between evolutionary replicates (same biotic context) against the difference between strains that evolved in a different biotic context. Differences are measured as the Euclidean distance between strains. Circles and error bars indicate the mean and the standard error of each pair (A) or species in a specific coculture (B).*

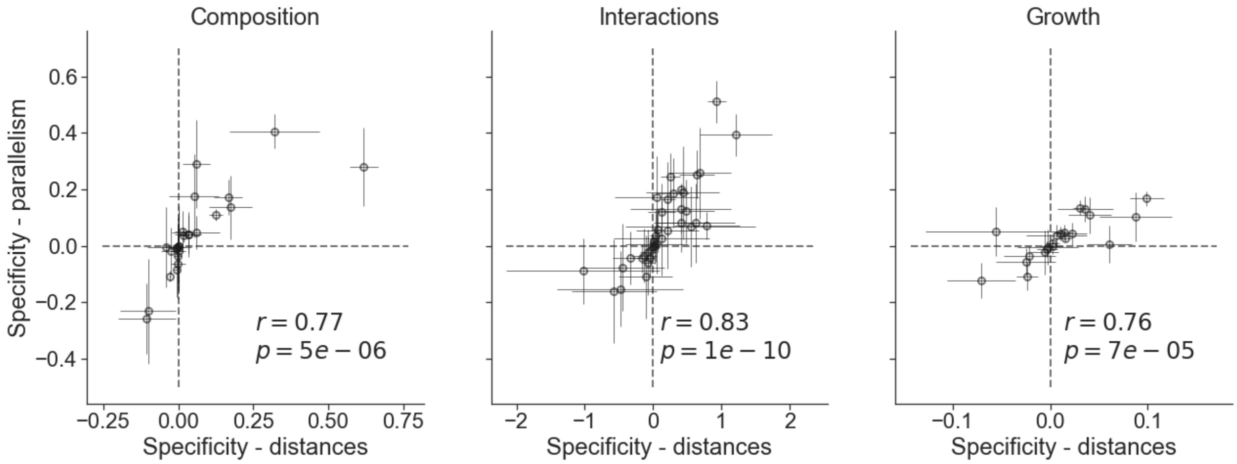


***Figure S24: Correlations between specificity scores calculated as distances and specificity scores calculated as parallelism.*** *Each panel represents all the scores calculated for each parameter. Statistics are the Pearson r and associated p-value for each panel.*

#### 6. Reproducibility

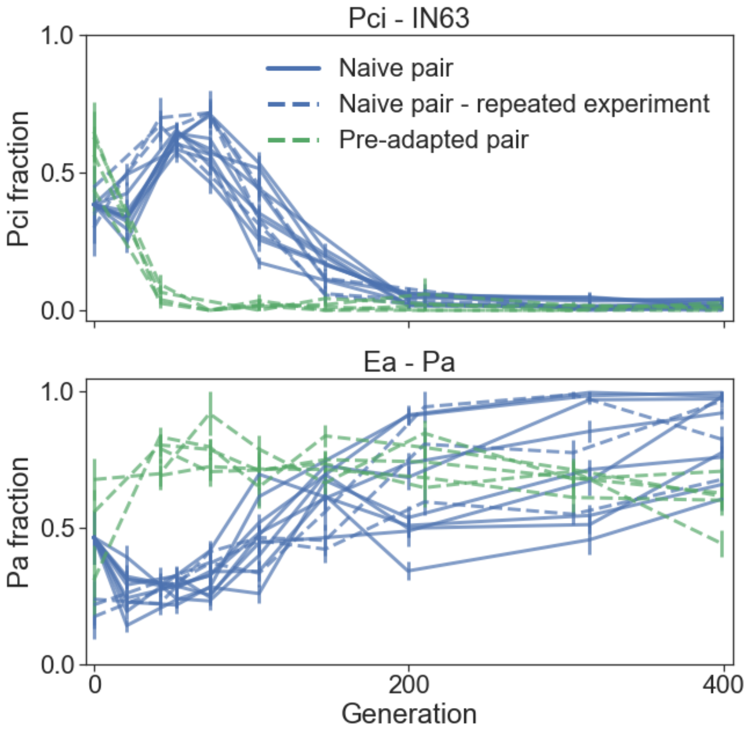


***Figure S25: Trajectories are similar across experiments, but not across evolutionary histories.*** *In order to verify that the two evolution experiments that were conducted are comparable, we included in the second experiment (Experiment Ev2) that focused on pre-adapted pairs also two naive pairs identical to those that were evolved in the first experiment (Experiment Ev1) . Each panel shows the fraction of one of the species in each pair throughout the evolution experiment, and lines represent different replicates. Blue full lines, and blue dashed lines are pairs assembled from the same stock in experiment Ev1 and Ev2 respectively. Green dashed lines are pairs assembled of pre-adapted strains, and were co-cultured in Experiment Ev2. Line style (full vs dashed) indicates the experiment, and line color indicates evolutionary history.*

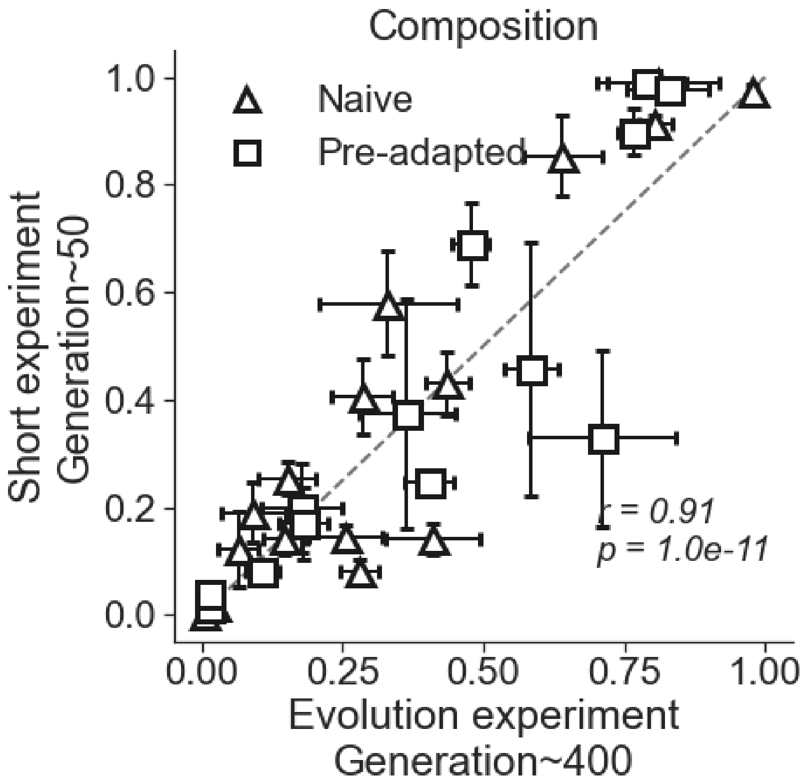


***Figure S26: Reproduction of composition data.*** *We compared the composition of pairs after ~400 generations of co-culturing (Experiments Ev1, Ev2), with their composition in a short ecological experiment after these pairs were frozen, thawed and co-cultured for ~50 generations (Experiment Ec1). Each marker and error bar represent the mean and standard error of the mean of each pair across replicates. Squares are pairs of naive strains (Experiment Ev1), and triangles are pairs of pre-adapted strains (Experiment Ev2). The purpose of this figure is to observe whether composition data is reproducible across experiments, however several non-technical mechanisms could cause differences between the experiments. For example if long ecological dynamics are required for stabilization, ~50 generations might not suffice. Similarly new mutations that shift the composition that arose at the end of the evolution experiment could affect the compositions in the short experiment. Nevertheless, composition of pairs after ~50 generations of re-inoculation and co-culturing was typically similar to the composition at the end of the experiments (Pearson r = 0.9, p-value = $10^{-11}$).*

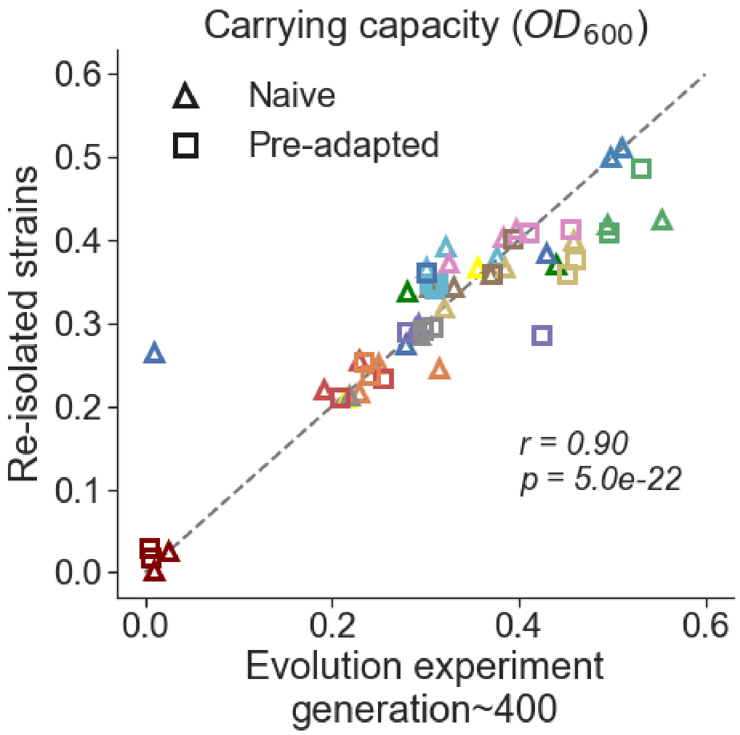


***Figure S27: Reproducibility of carrying capacity.*** *The optical density of strains that evolved in monoculture at the end of a 48h growth cycle at the 38$^{th}$ growth cycle (Experiments Ev1, Ev2), against their optical density at the end of the growth curve experiment (Experiment Gr1). Each market represents the optical density of a specific independently evolved strain. Triangles represent naive evolved monocultures (Experiment Ev1), and squares pre-adapted monocultures (Experiment Ev2). Pearson r = 0.9, p-value = $5*10^{-22}$.*

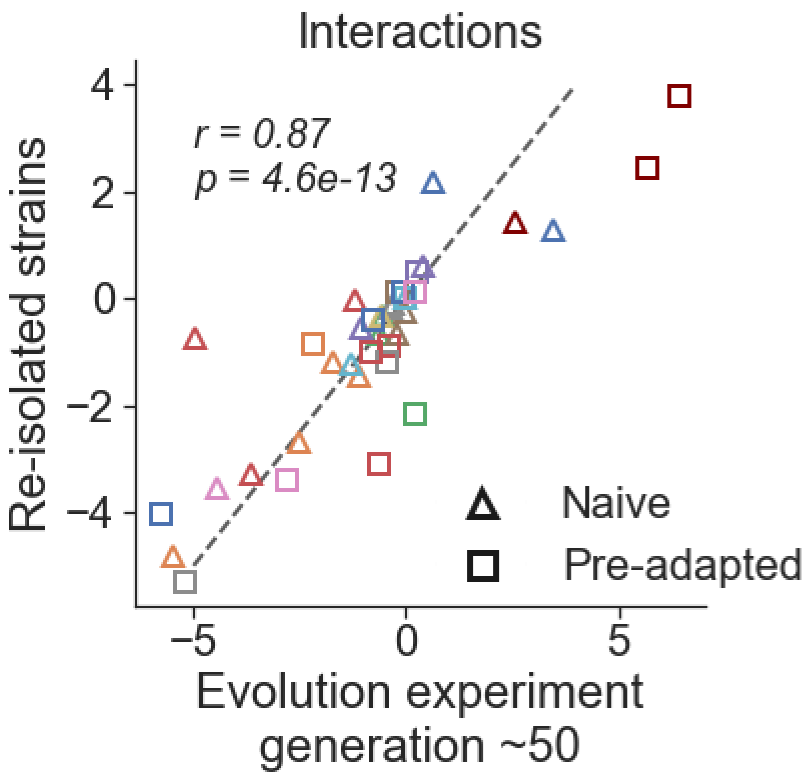


***Figure S28: Reproducibility of interactions:*** *Interactions calculated from the evolution experiments at generation ~50 (Experiment Ev1, Ev2), against the interaction of the ancestral strains in short experiments (Ec 2-5). Pearson r = 0.86, p =* $7*10^{-13}$.

#### 7. Mutations filtering

| Species | Gene name | Gene position | Number of mutated strains | Included in the analysis | Rationale for including/excluding |
|---|---|---|---|---|---|
| Ab | Ab_02713/nhaP | intergenic (-128/+345) | 2 | no | low coverage |
| | Ab_02767/rpoD | intergenic (-42/-325) | 2 | yes | |
| | ata | 768 | 8 | no | duplicated coverage |
| | | | 13 | no | duplicated coverage |
| Ea | Ea_02626/uxuA_2 | intergenic (-221/-40) | 3 | yes | |
| | Ea_04388 | 180 | 10 | no | duplicated coverage |
| | | 180 | 4 | no | duplicated coverage |
| | Ea_05065 | Multiple genes (1-3003) | 17 | no | Assembly issue |
| | Ea_05066 | Multiple genes (1-1493) | 17 | no | Assembly issue |
| | –/– | intergenic (–/–) | 17 | no | Assembly issue |
| | | | 17 | no | Assembly issue |

| | | | | | |
|---|---|---|---|---|---|
| Pa | Pa_02286/capB_2 | intergenic (+34/+40) | 3 | no | Two separate repetitive sequence elongations |
| | | intergenic (+52/+22) | 2 | no | Two separate repetitive sequence elongations |
| | Pa_03668–intA | Poly | 4 | no | low coverage in this region also in the ancestor |
| | Pa_04374/aceA | intergenic (+58/-442) | 2 | no | polyA elongation |
| | acs_2/Pa_05060 | intergenic (-30/-381) | 3 | yes | |
| | sasA_14 | coding (499/1314 nt) | 2 | yes | |
| Pch | dat_2/qseC_3 | intergenic (-246/+3) | 20 | no | In ancestor |
| | lutR_1 | coding (372-379/750 nt) | 2 | yes | |
| | proY_2 | coding (621-629/1422 nt) | 2 | yes | |
| | ptsP | coding (2045-2047/2280 nt) | 2 | yes | |
| | tesB/Pch_05630 | intergenic (-94/+7) | 20 | no | In ancestor |
| Pf | Pf_02401/Pf_02402 | intergenic (+338/+659) | 4 | no | In ancestor |

| | | | | | |
|---|---|---|---|---|---|
| | | intergenic (+345/+655) | 17 | no | In ancestor |
| | Pf_05002 | coding (853-855/2388 nt) | 5 | yes | |
| | aer_3/Pf_03656 | intergenic (+29/+86) | 8 | yes | |
| | alkB/Pf_01024 | intergenic (-148/+110) | 2 | yes | |
| | gatB/rlpA_1 | intergenic (+35/-21) | 8 | no | low coverage |
| | gcvT_2 | coding (743/1122 nt) | 2 | yes | |
| | glcB/glnG_2 | intergenic (+69/+11) | 14 | no | Shortening of a polyC sequence |
| | glnQ_1/mcpQ_1 | intergenic (+194/+822) | 2 | yes | |
| | norR_3 | 649 | 2 | yes | mutations occurred in separate areas of the gene, and same exact mutation occurred both in Naïve and preadapted |
| | puuA_1 | 46 | 5 | yes | multiple unique mutation, only two are identical, all non-synonymous |

| | | | | | |
|---|---|---|---|---|---|
| | puuA_1/puuA_2 | intergenic (-202/-466) | 3 | yes | multiple unique mutation, only two are identical |
| | spuD_1/spuE_1 | intergenic (+91/-71) | 2 | yes | |

***Table S7: Manually inspected mutations and reasoning that led to their exclusion.*** *"Low coverage" indicates that the number of reads mapping to the area of the mutations were consistently low; "duplicates coverage" indicates that the coverage of the mutated gene was at least 2 times higher than in its surrounding, suggesting that some variability could be possibly explained by multiple copies of the sequence; "Assembly issue" indicates that the mutation prediction seems to be related to error in the assembly process; "In ancestor" indicates that this mutation is predicted also when analyzing sequences generated from the ancestor against itself.*

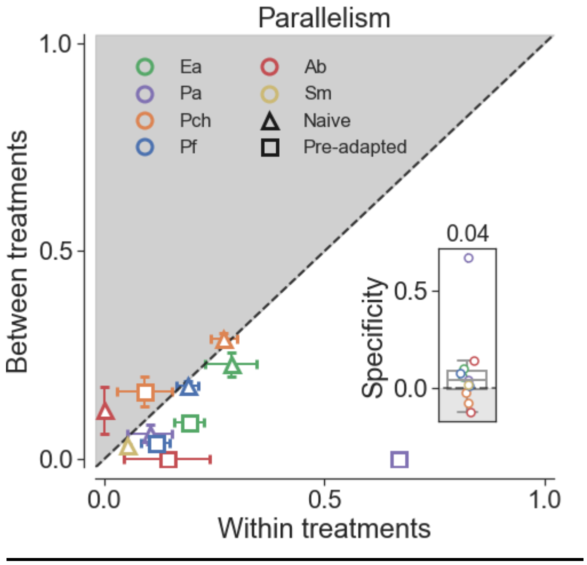


***Figure S29: Gene-level parallelism between and within treatments after excluding large deletions and mutations in intergenic regions.*** *Parallelism in genomic evolution is measured as the dice similarity in mutated genes. Error bars indicate the standard error mean for each species. Insets show the distribution specificity scores and the number above is the median score across all species. Triangles represent naive strains (evolved in Experiment Ev1), and squares are pre-adapted strains (evolved in Experiment Ev2).*

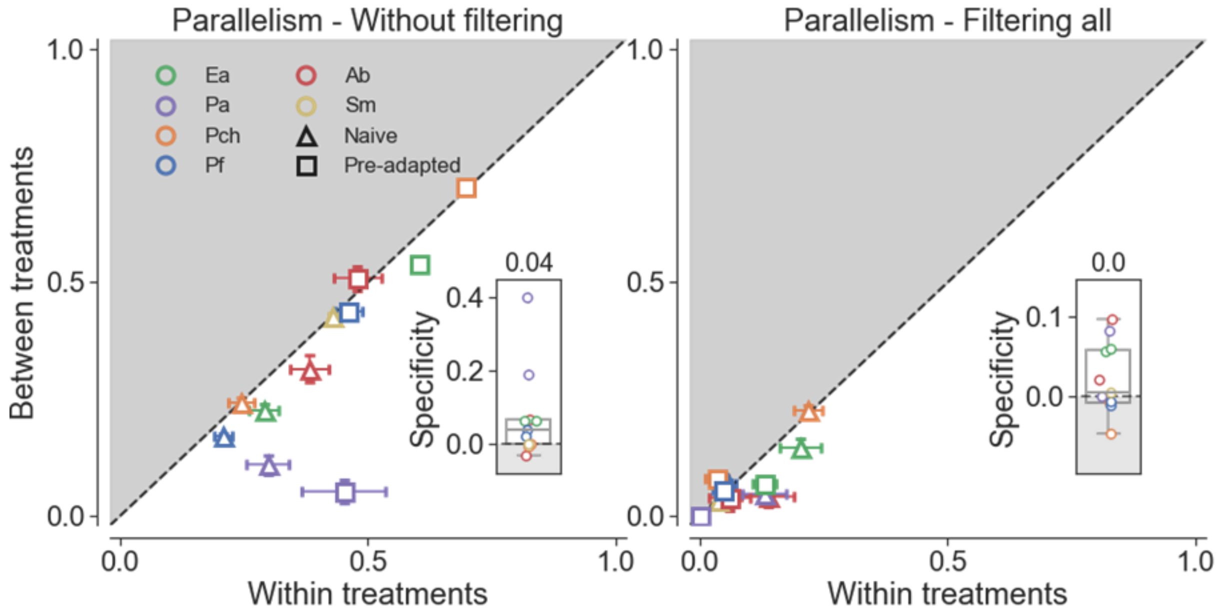


***Figure S30: Gene-level parallelism between and within treatments after different mutation filtering regimes.*** *(Right) without filtering parallelly mutated exact sequence mutations at all, (left) after filtering all parallelly mutated exact sequence mutations. Parallelism in genomic evolution is measured as the dice similarity in mutated genes. Error bars indicate the standard error mean for each species. Insets show the distribution specificity scores and the number above is the median score across all species. Triangles represent naive strains (evolved in Experiment Ev1), and squares are pre-adapted strains (evolved in Experiment Ev2).*
